# The genetic basis of adaptation to continuous environmental gradients informs predictions about climate change responses

**DOI:** 10.64898/2026.11.19.767859

**Authors:** Derek A. Denney, Emily B. Josephs, Megan L. DeMarche, Jill T. Anderson

**Author notes:** **Corresponding Author:** Jill T. Anderson, **Email:**.

## Abstract

A central goal of evolutionary genetics is to determine how environmental pressures maintain adaptive genetic variation and evaluate how this variation is encoded in the genome (1). Dissecting the genetic basis of local adaptation carries new urgency in an era of rapid climate change. We identified genomic regions associated with climatic adaptation, quantified the fitness consequences of allelic variation, and forecasted evolutionary responses to novel climates in the ecological model plant, *Boechera stricta*. We leveraged two complementary approaches: genome-environment associations (GEA) of climatic factors and genome-wide association studies (GWAS) of climate niche parameters derived from demographic models built from long-term field experiments. The GEA detected 154 unique SNPs linked to climatic variation. The GWAS identified 319 unique single nucleotide polymorphisms (SNPs) associated with the climatic niche. After pruning for linkage disequilibrium, we linked genotypic variation to fitness across broad climatic gradients. Genotypes at most loci showed non-linear seedling recruitment success in response to variation in snowmelt timing, with both conditional neutrality and genetic trade-offs contributing to local adaptation. Local adaptation via adult survival arose primarily through conditional neutrality. Candidate genes near these loci were implicated in stress responses. Contrary to expectations, increasing temperatures and accelerating snowmelt did not augment the absolute fitness of low elevation genotypes. Instead, our analyses revealed that the hot and dry conditions projected under climate change may rapidly exceed the tolerance of this species. Nevertheless, we identified 47 SNPs at which evolutionary rescue or assisted migration could partially ameliorate projected fitness declines under climate change.

**Significance Statement:** We extended genome-wide association approaches beyond traditional phenotypes to map the climate niche. Genotype-environment interactions for fitness across continuous climatic gradients revealed that polygenic local adaptation is driven by genetic trade-offs and conditional neutrality. Ascertainment bias could obscure cryptic genetic trade-offs if analyses consider only one fitness component or do not explore non-linear responses to environmental variation. In contrast with GWAS, genome-environment associations overemphasized locally maladaptive genetic variants, highlighting the limits of niche-agnostic approaches. Unexpectedly, cool temperatures with delayed snowmelt enhanced fitness for genotypes at nearly all SNPs detected through GWAS and GEA. Genetic variation relevant to climate change at 47 SNPs may reduce the risk of population contractions in response to novel climates.

## Introduction

Spatial variation in abiotic and biotic factors across the landscape has driven the evolution of local adaptation in taxa across the tree of life (2, 3). We know more about *the existence* of local adaptation than we do about the agents of selection that shape local adaptation (4, 5), or the genetic and genomic basis of local adaptation (6–8). This gap in our understanding of the mechanistic basis of local adaptation limits our ability to predict organismal responses to rapid climate change. By decoupling the conditions that characterized complex historical environments (9, 10), climate change has already disrupted local adaptation (11–14). The genetic basis and architecture of local adaptation strongly influence individual fitness and population persistence in projected future conditions (15, 16).

Local adaptation can evolve through genetic trade-offs (Figs. 1 and S1), in which local alleles have a fitness advantage relative to foreign alleles (e.g., 17). Alternatively, the local allele may have a fitness advantage in the home climate without experiencing a fitness cost at the opposite end of the climatic gradient (i.e., conditional neutrality, 8, 18). Conditional neutrality (Fig. 1B) can only contribute to local adaptation if multiple polymorphic loci show contrasting patterns (6, 18), as is the case when a high latitude allele is conditionally beneficial at high latitudes at one quantitative trait locus (QTL) and a low latitude allele is conditionally beneficial at low latitudes at an unlinked locus in *Boechera stricta* (Brassicaceae) (19). With sufficient gene flow, the conditionally advantageous allele would spread across the landscape; thus, conditional neutrality may represent a transient phase in the evolution of local adaptation (18, 20).

**Figure 1.**
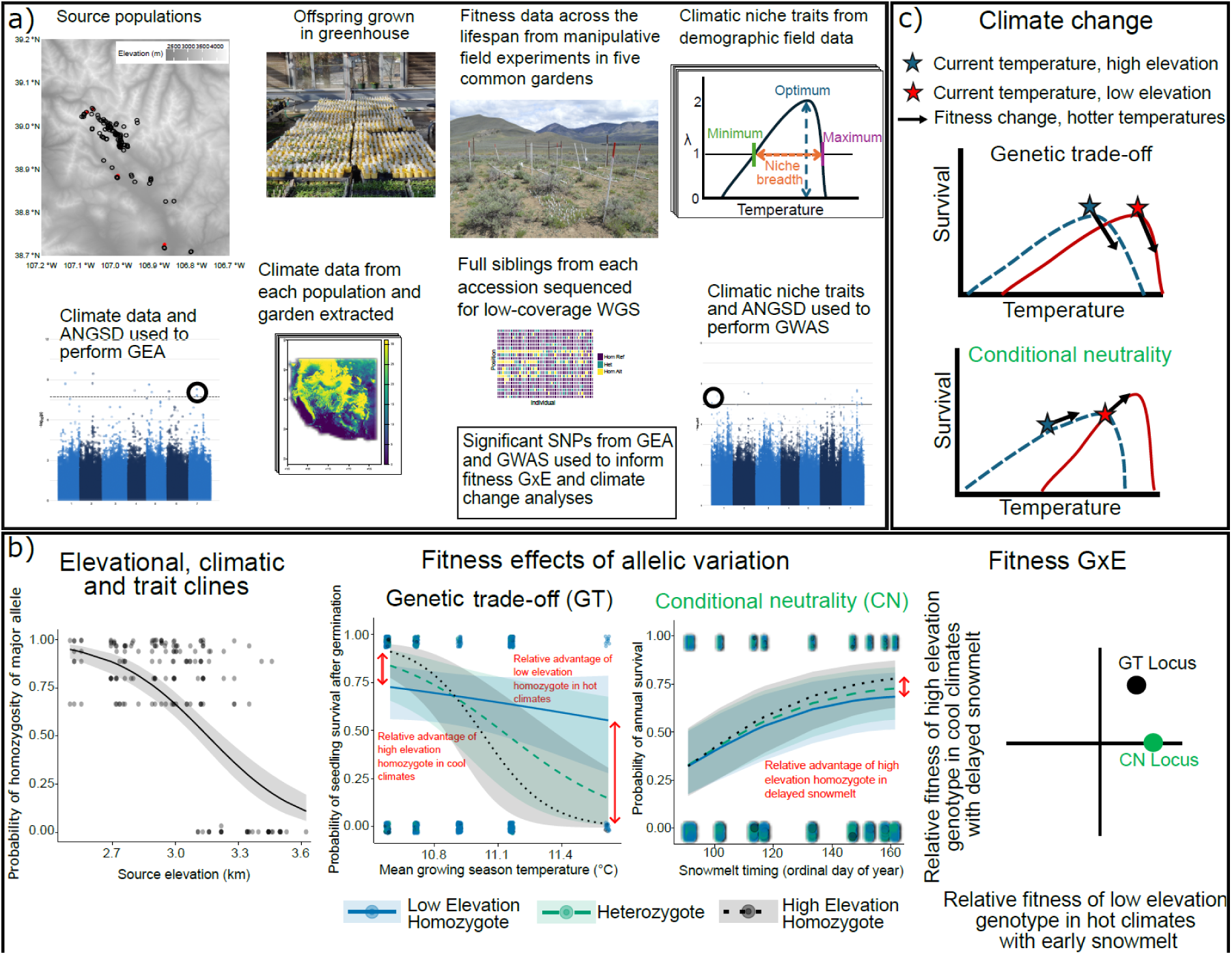
Project workflow. (A) Seeds were collected from source populations, grown in the greenhouse to homogenize maternal effects, and transplanted into 5 common gardens (red circles on the map) in which we manipulated snowmelt timing and temperature. Fitness data from the field experiment were used to model the climatic niche (14). One full sibling per accession was sequenced using low coverage whole genome sequencing (WGS). We conducted genome-environment associations (GEA) using climatic data for each source population and genome-wide associations (GWAS) using climate niche parameters for each accession. (B) Significant SNPs were used to examine elevation, climatic and trait clines in the probability of homozygosity of the major allele (e.g., SNP 1_6920936, Fig. S6), and then evaluate the contributions of genetic trade-offs (e.g., SNP 4_13558274) and conditional neutrality (e.g., SNP 1_14098322) to local adaptation and test the fitness consequences of climate change. We demonstrate where these two loci would be located on a figure highlighting the relative fitness of the local allele in the home climate (see Fig. 3A and B). (C) Variation in the thermal niche could alter fitness under elevated temperatures for loci displaying genetic trade-offs vs. conditional neutrality.

Reciprocal transplant experiments have evaluated the genetic basis of local adaptation in response to discrete environments (21), which cannot capture non-linear fitness functions that arise across continuous environmental gradients. Species often display asymmetrical thermal performance curves, with steep declines in fitness at elevated temperatures (Fig. 1) (22). Accessions originating from disparate climates likely evolve distinct niche parameters, such as climatic optima (14) and niche breadth, both of which could influence the extent of local adaptation and the degree to which novel climates alter fitness (Fig. 1C). If local adaptation arises from loci displaying genetic trade-offs and current climates optimize fitness, higher temperatures could depress the fitness of local alleles across the landscape (Fig. 1C), but gene flow from lower latitude or elevation populations could introduce warm-adapted alleles into local populations (16, 23, 24). If instead, the niche breadth is wide or genotypes currently inhabit suboptimal climates, conditional neutrality could cause fitness to increase as the climate warms (Fig. 1C). The fitness responses to novel climates are likely dictated by the shape of the climatic niche, current and future climatic conditions, and the genetic basis of local adaptation, with outcomes potentially differing across loci that display conditional neutrality vs. genetic trade-offs.

Rigorous tests of the mechanistic basis of local adaptation require the manipulation of putative agents of selection (25), ideally in field experiments within the native environment in which sequenced lineages evolved. To that end, we conducted a multi-year field experiment of the widespread ecological model species, *B. stricta*, in which we manipulated snow dynamics and growing season temperature in five common gardens arrayed across an elevational gradient in the Elk Mountains (Colorado, U.S.A.) (14). A previous study leveraged these data to model accession-specific thermal and snowmelt niches in a climate-driven demographic model (14). To identify regions of the genome associated with climatic adaptation, here we first linked climatic factors to sequence variation using genome-environment association (GEA) in analyses agnostic to fitness variation (N=95 populations, Figs. S2-S4) (7). We then conducted a genome-wide association study (GWAS, 219 accessions from 115 source populations; Table S1) of 10 climate niche parameters derived from the demographic models (14), including the timing of snowmelt and the growing season temperature that optimize lifetime fitness (per capita rate of increase, λ). Finally, using fitness and sequence data from the common gardens (N=14,170 adult plants and 219 accessions; and N=33,211 seeds and 191 accessions, SI Methods), we: (a) evaluated the contributions of conditional neutrality and genetic trade-offs to the genetic basis of climatic adaptation at SNPs detected through GEA and GWAS; (b) examined whether multivariate climate change is predicted to alter fitness of genotypes at SNPs implicated in climatic adaptation; and (c) identified candidate genes with allelic variation that could facilitate adaptation to novel climates. We validated our local adaptation results with data from a smaller reciprocal transplant experiment (N= 5,909 transplants and 97 accessions, SI Methods).

## Results and Discussion

### Genetic architecture of local adaptation

After quality filtering, 2,737,409 loci were retained (SI Methods). Our GEA detected 1165 SNPs associated with ecologically-relevant climatic factors and the GWAS identified 366 SNPs associated with the thermal and snowmelt niche (Table S2; Fig. 2 and S5). After collapsing duplicate SNPs across phenotypes, we recovered a total of 473 unique SNPs, of which 319 were exclusive to the GWAS and 154 were unique to the GEA. Overall, our analyses implicate genes involved in signaling, stress response, and transport processes (Table S3). Whether local adaptation arises through few linked loci of large effect clustered in specific regions of the genome or many loci of small effect distributed throughout the genome depends on the spatial extent of gene flow, the magnitude of divergent selection, and the degree of genetic drift (26, 27). The large number of loci identified across GWAS and GEA indicates that climatic adaptation in *B. stricta* is polygenic, consistent with theoretical expectations for a self-pollinating species with limited gene flow (27). Nevertheless, adaptation is not so polygenic as to preclude identifying alleles with clinal patterns (Fig. 1 and S6). Spatial genomic simulations suggest that, depending on the genomic architecture, high polygenicity can slow adaptation and increase maladaptive responses to climate change (16), corresponding with demographic models predicting population declines under climate change in *B. stricta* especially at the trailing edge of the range (14).

**Figure 2.**
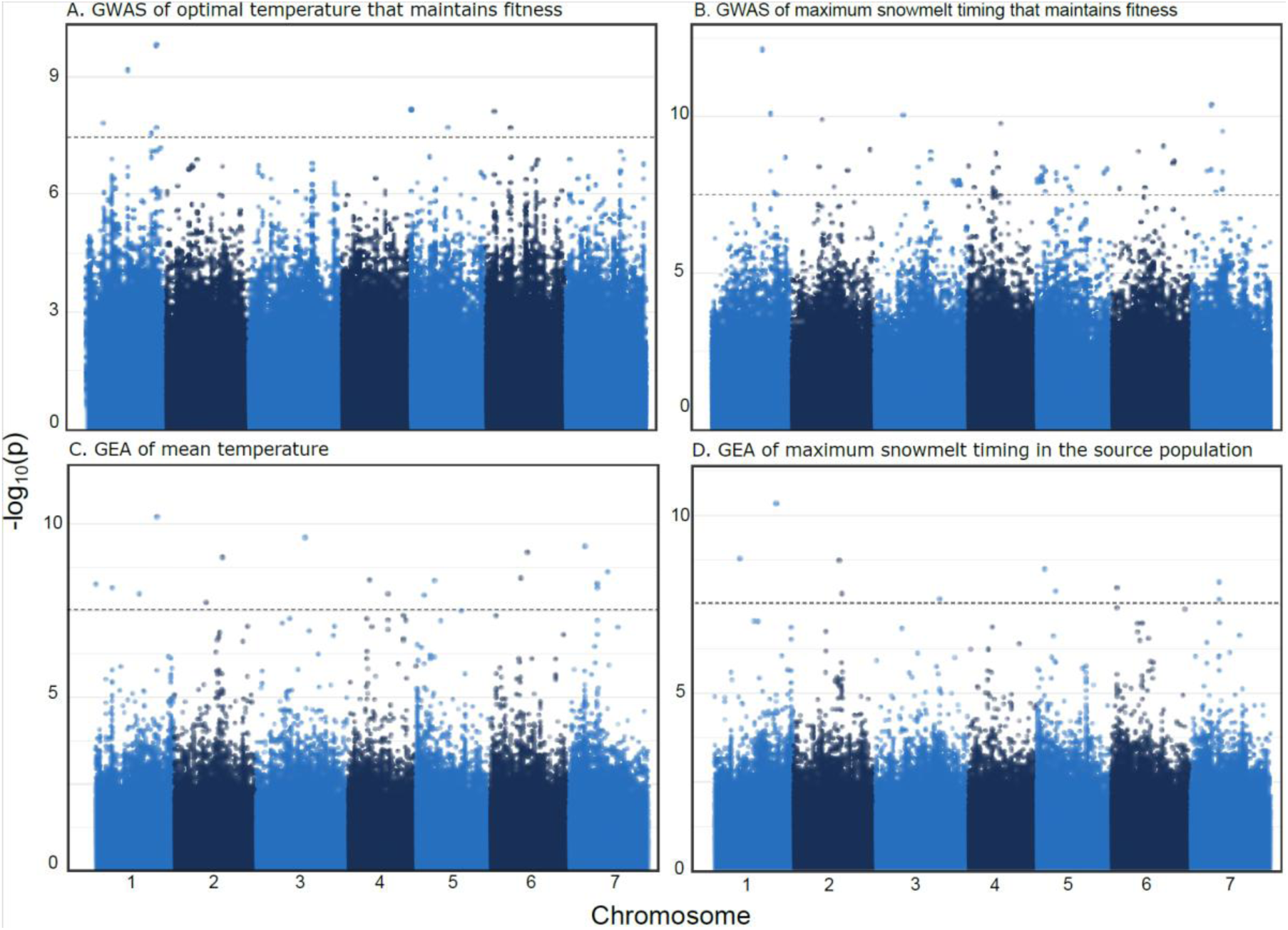
Representative Manhattan plots displaying results of genome wide association for (A) optimal temperature and (B) maximum snowmelt timing beyond which fitness declines to unsustainable levels, and genome-environment association for (C) mean temperature, and (D) maximum snowmelt timing in the source population. Fig. S5 shows all Manhattan plots.

### Genetic basis of local adaptation

After pruning for linkage disequilibrium and analyzing clinal variation, we identified 41 GWAS loci and 40 GEA loci that showed significant elevational clines in major allele homozygosity (Fig. 1 and S6; Table S4-S7), enabling us to determine the elevational origin of major and minor alleles. We predicted that low elevation genotypes would have enhanced fitness in hot, dry climates with early snowmelt whereas high elevation alleles would show the greatest fitness in cool, mesic climates with delayed snowmelt (Fig. S1), consistent with climatic gradients in this region (Figs. S2-S4). We assessed fitness components that influence long-term population growth rates in perennial species (28) including *B. stricta* (14): seedling recruitment and annual adult survival.

Genotype by climate interactions significantly influenced fitness in 48 of 105 analyses of annual adult survival and 37 of 105 analyses of seedling recruitment, after correction for multiple testing (29). In linear models of adult survival, 2 GWAS SNPs and 4 GEA SNPs exhibited genetic trade-offs (Table S8; Figs. 1 and 3). Genetic trade-offs were more evident for seedling recruitment for which we found negative quadratic fitness functions for 20 SNPs in response to the timing of snowmelt. At 15 GWAS SNPs and 2 GEA SNPs, the low elevation allele displayed optimal seedling fitness under earlier snowmelt than the high elevation allele (Table S8; Fig. 4), concordant with predictions. Whether we interpret a SNP as displaying conditional neutrality or genetic trade-offs depends on the portion of the climatic space examined, highlighting the importance of considering non-linear genotypic variation in fitness across complex climatic gradients.

**Figure 3.**
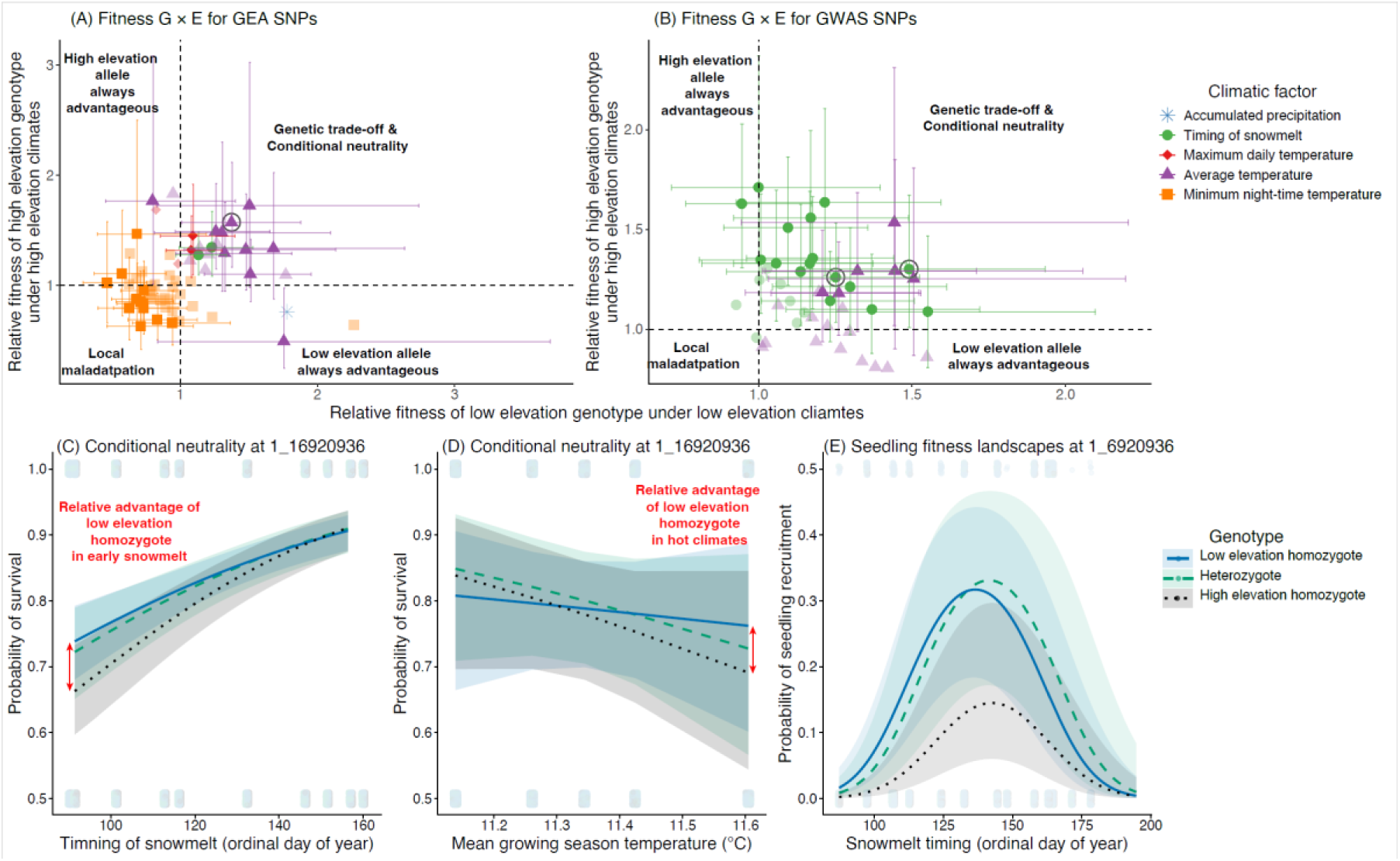
Genetic basis of local adaptation for the set of SNPs from (A) GWAS and (B) GEA. The data points reflect the odds ratios ± 95% confidence intervals (CI; error bars) of survival for the low vs. high elevation genotype in hot and dry environments with early snowmelt (X-axis) and the high vs. low elevation genotype in cool, moist climates with delayed snowmelt (Y-axis). Odds ratios >1 indicate that the local genotype outperforms the foreign genotype in the home climate. Data points are color coded by the climatic factor included in the fitness model, and symbols with greater transparency and no error bars had non-significant genotype by climate interactions after correction for multiple testing. Genetic trade-offs (circled loci) are present if the odds ratios (OR) and 95% CI are fully within the upper righthand quadrant of each panel. Conditionally neutral loci have 95% CI bars that span OR = 1 in one climate but in the contrasting climate. Local maladaptation was only present in GEA SNPs. (C) Conditional neutrality in response to snowmelt timing and (D) temperature at a representative SNP (1_16920936). (E) Genetic trade-offs were more apparent at 1_16920936 for seedling recruitment when examining the earlier peak snowmelt timing for the low elevation homozygote than the heterozygote; both the low elevation homozygote and the heterozygote outperformed the high elevation homozygote across most snowmelt timings.

**Figure 4.**
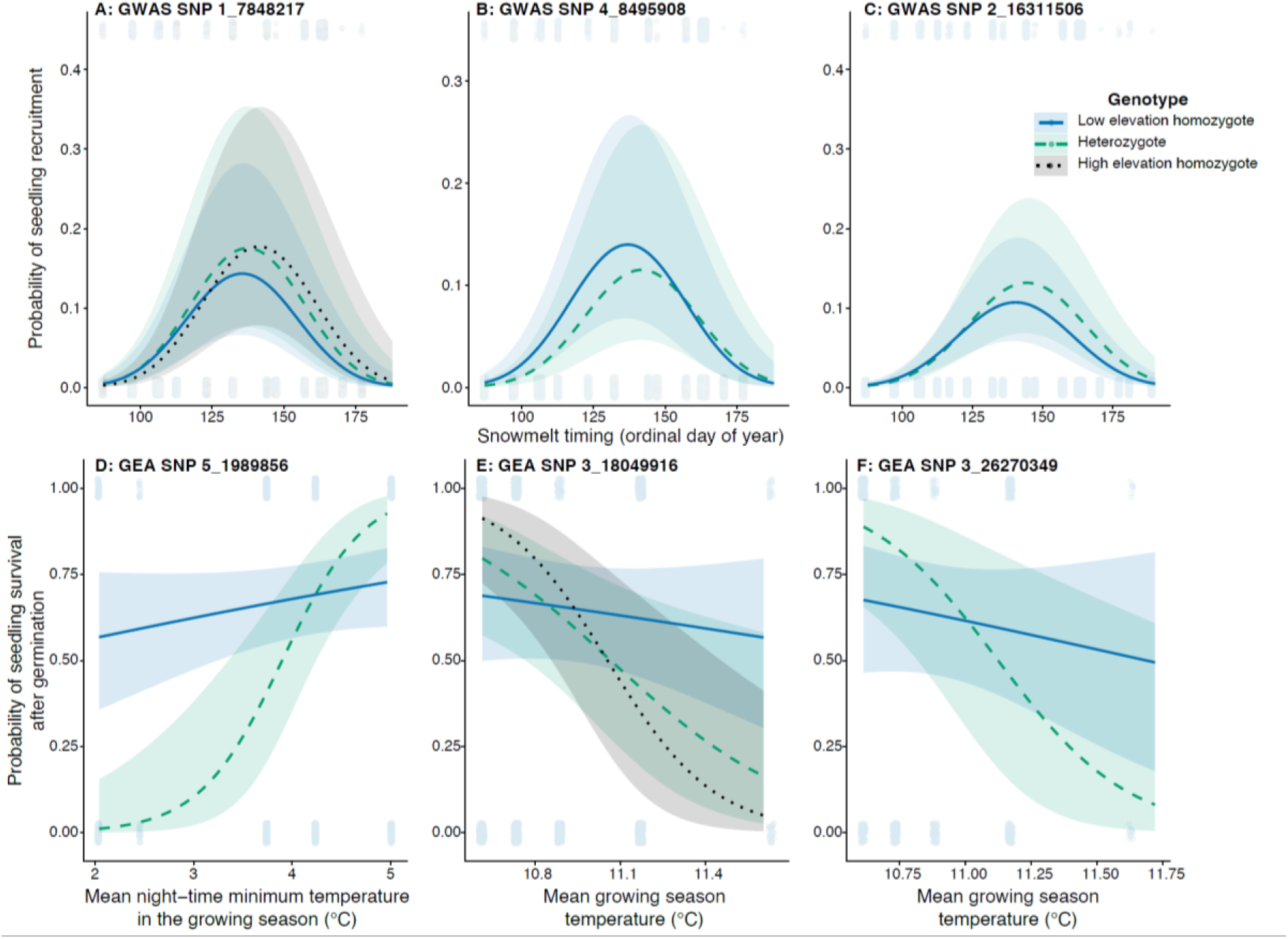
Complex fitness landscapes for seedling recruitment, with the low elevation genotypes displaying earlier optimal snowmelt timings than high elevation genotypes at most SNPs, including: (A) 1_7848217 (genetic trade-offs when considering the high elevation homozygote and the heterozygote, with conditional neutrality when examining the high vs. low elevation homozygotes); (B) 4_8495908 (conditional neutrality with low elevation homozygote maintaining higher fitness than the heterozygote at early snowmelt timings); and (C) 2_16311506: (conditional neutrality with the heterozygote displaying higher fitness than the low elevation homozygote at delayed snowmelt timings). For SNPs mapped to temperature variables in the GEA, we found (D) local maladaptation (e.g.,SNP 5_1989856), (E) genetic tradeoffs (e.g., SNP 3_18049916), and (F) conditional neutrality (e.g., 3_26270349).

We detected conditional neutrality at 24 SNPs (Table S8; Fig 3). More evidence for conditional neutrality emerged in the GWAS (15 SNPs) than the GEA (9 SNPs). Theory predicts that GEA is best suited to the detection of genetic trade-offs and lacks the statistical power to detect conditional neutrality (20), yet our GEA found more SNPs showing conditional fitness effects than genetic trade-offs. Ascertainment bias can restrict the discovery of genetic trade-offs, as local alleles must show a fitness advantage in at least two environments (18, 30) or (in our study) at opposite ends of the climatic spectrum. Indeed, our analyses of annual survival recovered crossing reaction norms for fitness for 9 GWAS and 7 GEA SNPs for which the relative fitness of the local vs. foreign genotype at one end of the climatic gradient failed to achieve statistical significance, causing us to render a determination of conditional neutrality (Table S8; Fig. 1 and 4F; Figs. S7-S8). Thus, genetic trade-offs remain challenging to identify even in long-term studies with experimental manipulations.

The fitness landscape often shifted *across* fitness components or experiments for the same SNP. For example, we discovered genetic trade-offs or conditional neutrality for annual survival and non-linear fitness functions reflecting local adaptation for seedling recruitment at 14 GWAS and 2 GEA SNPs (Table S8; Fig. 3C-E), illustrating that genetic trade-offs could be mistaken for conditional neutrality if only one fitness component is measured or if early life history stages are neglected. Results from a smaller reciprocal transplant experiment used for validation broadly align with the main experiment, showing significant climate by genotype interactions at 6 SNPs and marginally significant interactions at 6 additional SNPs, most of which demonstrated conditional neutrality (Table S9). The directionality of conditional neutrality shifted between the two experiments for four SNPs, which could signify that genetic trade-offs underlie local adaptation when the two datasets are considered together. For example, in the main dataset, the low elevation allele at SNP 1_1701580 was conditionally advantageous under early snowmelt, whereas in the validation dataset, the high elevation allele was conditionally beneficial in delayed snowmelt (Tables S8 and S9; Fig. S9). Thus, findings of conditional neutrality could reflect genetic trade-offs that remain cryptic (18) if ontogenetic stages differ in their response to environmental variation (31), if agents of selection differ in their allele-specific fitness effects, if insufficient environmental variation has been sampled, or if the true agent of selection was not investigated.

Comparative studies have consistently identified a set of core functional pathways underlying climatic adaptation in plants, including phenology, hormone signaling, and abiotic stress responses (32). In our study, the candidate loci for SNPs associated with local adaptation often overlapped with these pathways at the broad functional levels, especially for genes involved in signaling, ion transport, and abiotic stress responses (Table S10). For example, a probable 2-oxoglutarate-dependent dioxygenase (AOP1 homolog; SAD12v2.2_chr1_gene1763) is part of the glucosinolate biosynthetic pathway, which is important for defense and environmental interactions (33). This finding is consistent with experimental evidence linking secondary metabolism and climate-mediated selection in *B. stricta* (34). Near multiple candidate SNPs, we also identified a U-box domain-containing protein (PUB5 homolog; SAD12v2.2_chr1_gene1555), which helps regulate protein turnover during drought and other abiotic stresses (35, 36). Other candidate genes spanned hormone and stress-related metabolism, vesicle trafficking, amino acid metabolism, and core cellular processes (Table S10). We could not identify candidate genes for eleven intragenic SNPs. Expansion beyond traditional genetic model systems can reveal new loci associated with in local adaptation.

#### Maladaptation

We found: (a) reciprocal local maladaptation, wherein local alleles are disfavored at both ends of the climatic gradient (5 SNPs), and (b) conditional maladaptation, in which the local allele has a fitness disadvantage at only one end of the climatic gradient (15 SNPs; Table S8). Locally maladapted quantitative trait loci (QTLs) have also been detected in *Arabidopsis thaliana* (8), and multiple loci with conditionally deleterious mutations could theoretically contribute to organismal local adaptation (18). Maladaptation in our system emerged exclusively for SNPs identified via GEA in response to average minimum temperatures during the growing season. Given that night-time temperatures have warmed at a greater rate than day-time temperatures (37), this maladaptation could reflect a biological response to a climate change factor that is often neglected in the eco-evolutionary literature.

#### Climatic agents of local adaptation

Initial GWAS and GEA results suggest adaptation is structured primarily along thermal gradients. The GWAS signal was highly skewed toward thermal niche parameters, which accounted for 82.5% of all hits (302/366), whereas snowmelt niche traits yielded substantially fewer associations (Table S2). In the GEA, SNPs were primarily associated with temperature and dewpoint variables (Table S2). Although GEA holds great promise for identifying environmental factors associated with local adaptation, genomic associations may reflect suites of correlated climatic factors rather than the true selective agent. Indeed, we detected more evidence for local adaptation and conditional neutrality in response to the timing of snowmelt (19 SNPs) than to temperature variables (16 SNPs; Table S8), despite the much larger number of temperature associations (1366 total) than the snowmelt associations (111 total) in the GEA and GWAS. This disconnect highlights the necessity of leveraging system-specific understanding of agents of selection, along with testing the fitness effects of genetic variation at SNPs identified via this approach.

Finally, the GEA identified more than three times the number of snow-associated SNPs using snowmelt variables calculated for local populations (14) from long-term Landsat 5-9 observations of normalized differential snow index (38) than it did with variables extracted from the SnowClim product (39). Thus, fine-grained local climatic data can prove more powerful than high-resolution gridded climate databases because they better match the selective environment experienced by the individual (40).

### Climate change

We expected fitness to increase with elevated temperature and accelerated snowmelt for low elevation alleles, as these are the conditions associated with low elevation environments (Fig. S1-S4); if true, low elevation populations may harbor genetic variation that could augment fitness under projected climates. For all SNPs displaying genetic trade-offs and for many conditionally neutral SNPs, the low elevation genotype outperformed the high elevation genotype in hot and dry environments with early snowmelt (Tables S8 and S9), consistent with the local vs. foreign criterion of local adaptation (2). Despite this *relative* fitness advantage at these SNPs, and inconsistent with the home vs. away criterion of local adaptation (2) the absolute fitness of the low elevation genotype either declined with increasing temperature and earlier snowmelt timing (17 SNPs, Figs. 3 & 4) or the slope of the relationship was non-significant (12 SNPs, Table S10). This pattern raises concerns about the fitness consequences of climate change.

Even when projected climates could depress adult survival of all genotypes at a SNP, the slope of the decline was often shallower for the low elevation than the high elevation allele, and genotypic reaction norms typically crossed at snowmelt timings and temperatures currently associated with low (2710 m) or mid-elevation (2890 m) locations (Fig. 3D, Figs. S7-S10). Mid-elevation climates represent a crossover point for seedling recruitment, as well, with low elevation alleles only providing a fitness advantage in warmer and drier climates (e.g., Fig. 4). The efficacy of assisted migration programs (41) may depend, in part, on the genetic basis and genomic architecture of local adaptation and the location in climate space where allelic reaction norms cross, as gene flow could inadvertently introduce maladapted alleles into distant recipient populations (16).

Our local adaptation analyses focused exclusively on SNPs with significant elevational clines. To generate more robust predictions of climate change responses, we examined climate by genotype interactions for fitness with the full set of 205 SNPs retained after pruning for linkage disequilibrium. Additionally, in these analyses we included transplants exposed to elevated temperatures in open-top chambers, which we excluded from the local adaptation analyses as the increase in temperature of 4.1° C far exceeded the range of spatio-temporal variation at these sites (14). Global change is simultaneously altering multiple abiotic conditions (42), which can interact in complex ways to generate increasingly novel climates (43) that shift the adaptive landscape. Thus, unlike the local adaptation analyses, we examined fitness as a function of all two- and three-way interactions of SNP genotype, temperature, and snowmelt timing to generate predictions under projected end-of-century elevated temperatures and novel snow dynamics.

Consistent with predictions of population declines for *B. stricta* (14), projected future temperatures depressed fitness for all genotypes at 112 SNPs (main effect of temperature; Fig. 5 and Tables S11 and 12). We found significant genotype by temperature interactions at an additional 28 SNPs (all from GWAS), revealing variation in the rate of decline of fitness with increasing temperature (14 SNPs), or no significant effect of temperature variation on fitness for one genotype (14 SNPs). Thus, although the predominant effect of increasing temperature was a reduction in fitness, for 14 SNPs, at which one genotype could maintain fitness under elevated temperatures.

**Figure 5.**
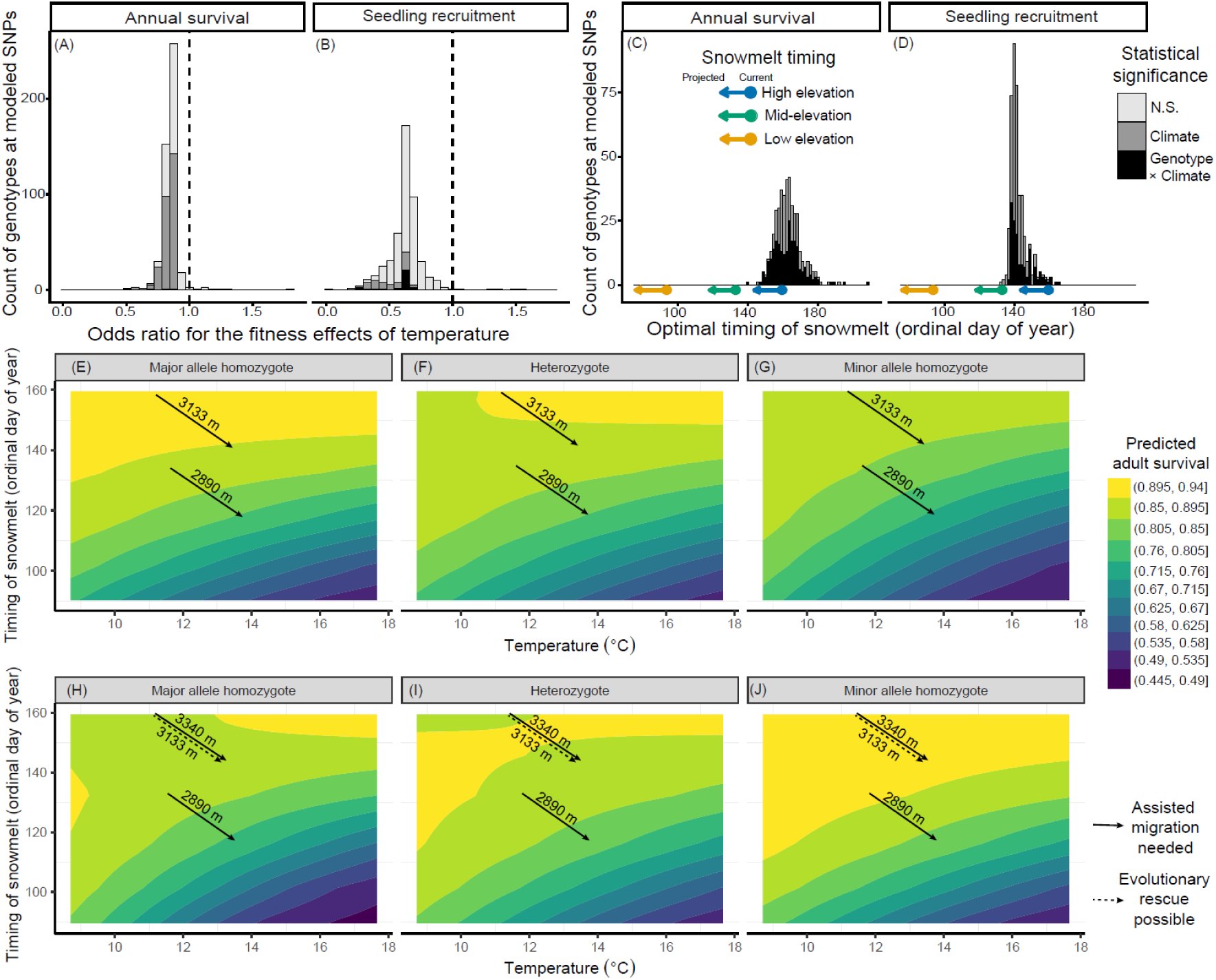
Fitness consequences of genotypic variation at 205 loci uncovered via GWAS and GEA and predictions of climate change responses. Histograms of odds ratios (OR) for (A) adult survival and (B) seedling recruitment, with OR <1 (dashed vertical lines) indicating that fitness declined with increasing temperature. Optimal snowmelt timing for (C) adult survival across 201 SNPs and (D) seedling recruitment across 205 SNPs. The current snowmelt timing (circle) and end-of-century projected snowmelt timing under RCP4.5 (end of arrow) for high (3340m, blue), mid-(2890m, green), and low (2553m, orange) elevations. For panels A-D, stacked histogram indicates whether models recovered a climate by genotype interaction (black), a main effect of climate (dark grey) or no significant effect of climate. Contour plots for adult survival in response to variation in temperature and snowmelt timing for all genotypes at SNPs (E-G) 5_11125224 and (H-J) 6_1311701. Arrows show current (start) and end-of-century projected climates at RCP 4.5 (arrowhead) for garden locations for which predicted future fitness differs across genotypes. For 5_11125224, the major allele homozygote has higher predicted fitness than the other genotypes under future climates at mid-elevation locations, where local populations are currently fixed for the minor allele (Table S14). For 6_1311701, the minor allele homozygote outperforms the other genotypes in projected future climate scenarios in three sites (2890m, 3133m, and 3340m). All three genotypes are present at 3133m (where evolutionary rescue is possible), but the populations at 2890m and 3340m are fixed for the major allele, requiring assisted migration.

Snow-dominated ecosystems at high elevations and latitudes have experienced diminished snowpack and accelerated snowmelt timing (44, 45). Advancing snowmelt significantly reduced annual survival for all genotypes at four SNPs, with G × E at two loci revealing genotypic variation in the extent to which projected snowmelt timings reduced survival (Tables S11). For the remaining SNPs (201 SNPs for survival and 205 for seedling recruitment; Table S11 and 13), we recovered significantly negative quadratic fitness functions, enabling us to estimate the snowmelt timing at which each genotype displays the highest fitness. Across genotypes, the average optimal snowmelt timing was June 1 in a non-leap year (ordinal day 152.6, range: 131.9 - 208.5; Table S13; Fig. 5), reflecting current and projected climates at the two higher elevation gardens (Table S14). The optima at these loci are already later than current snowmelt timings at the two lowest elevation gardens, concordant with local maladaptation at the organismal level in low elevation *B. stricta* populations (14) and with broad-scale population contractions at the trailing-edge of the ranges across species (31, 46).

We cross referenced SNPs with significant genotype by climate interactions against genotypic frequencies at these loci in the five natural populations adjacent to each common garden (Table S14-15), identifying 47 SNPs for which one genotype has the highest predicted fitness under projected climates in at least one location. We classified 40 SNPs as potential *evolutionary rescue loci* because the allele with the highest predicted future fitness currently exists in low frequency in at least one local population (Table S14; Figure 5H-J), suggesting that adaptive evolution via standing genetic variation could minimize fitness declines in response to earlier snowmelt and increasing temperatures. We characterized an additional 4 SNPs as potential *assisted migration loci* because the allele with the highest predicted future fitness does not currently exist in the location where it will be favored, but it does occur in up- or downslope populations (Table S14; Figure 5E-J). Finally, 3 SNPs are either *evolutionary rescue* or *assisted migration* loci, depending on the context (Table S14). These designations do not apply evenly across all five natural populations. Instead, we found the greatest potential for evolutionary rescue or assisted migration for genotypes homozygous for the major allele at our second to highest elevation natural population (3133m above sea level). We only found evidence for genotypic variation in projected future fitness in the three lower elevation gardens at 9 SNPs, suggesting limited evolutionary potential in low elevation sites.

Assisted migration (41) programs that consider only upslope translocations may miss opportunities to spread beneficial alleles across the landscape. Climate change exerts strong selection on traits (47), which could favor novel suites of phenotypes, such as the early flowering of high elevation accessions and the drought tolerance and herbivore resistance of low elevation populations (48, 49). For three loci in our study, downslope gene flow could redistribute alleles adaptive under future climates. Furthermore, at SNPs 2_10718142 and 5_2787751, the heterozygote has the highest predicted fitness in future climates at the lowest and mid-elevation sites (Table S14). Assisted gene flow and subsequent introgression, even in a species that primarily self-pollinates, could introduce alleles that may be better able to withstand environmental change.

Candidate genes near the evolutionary rescue and assisted migration SNPs (Table S16) included homologs of *Arabidopsis* genes with roles in abiotic stress responses and root functions, such as homologs of DEAR4 (SAD12v2.2_chr7_gene4955), a transcription factor that promotes heat tolerance through induction of heat shock proteins (50), and LTI65 (SAD12v2.2_chr7_gene3663), which is induced by cold and drought stress (51). We also identified candidate homologs of PRX66 (SAD12v2.2_chr3_gene4481), a peroxidase involved (52) in drought stress response (53), and a homolog of FRO2 (SAD12v2.2_chr1_gene65), which has been implicated in drought and thermal stress responses (54, 55).

## Conclusions

We detected hundreds of SNPs that could be associated with climatic adaptation in *B. stricta* through a combination of GWAS using climate niche parameters and GEA of climatic factors, highlighting the polygenic nature of climatic adaptation. Genetic trade-offs and conditional neutrality contribute to local adaptation in this system, and non-linear fitness landscapes blur the distinction between these mechanisms. Genome-environment association approaches are increasingly accessible because climatic data can easily be downloaded from publicly available repositories (7); however, this approach failed to detect key SNPs that are statistically associated with local adaptation to climate in this system and identified 20 locally maladapted SNPs. GEA may not reveal the true genetic basis of climatic adaptation. Climate change is likely pushing natural accessions beyond their climate niche, as accelerated snowmelt and elevated temperatures depressed survival and seedling recruitment at most climate-related SNPs. Up- and downslope assisted migration could redistribute genetic variation that could forestall population declines. Standing genetic variation within populations at 43 loci could lessen reductions in fitness under projected climate change, but that variation may not be enough to counteract fitness declines that could arise in response to novel climates.

## Materials and Methods

### Study System

*Boechera stricta* (Graham) Al-Shehbaz (Brassicaceae) is a perennial forb native to North America from New Mexico through Alaska, at elevations 700-3900m above sea level (56). This species primarily self-pollinates (57) and has limited seed dispersal, restricting gene flow (14) and leading to high homozygosity and low within-population quantitative genetic variation in fitness-related traits (58, 59). In Rocky Mountain meadows, temperatures, growing season length and aridity decline with elevation and snow melts later in the season (60) (Fig. S2-4). Mountainous ecosystems are subject to high rates of warming and extreme loss of snow cover (61). In the western U.S., temperatures have increased, snow accumulation has declined, and the timing of snowmelt has advanced since the mid-1900s (44, 62). Models suggest Colorado could lose 50-60% of its snow cover by 2080 (63), as may many regions in the western U.S. (45). Decreased snowpack and heightened evapotranspiration from hotter temperatures augment aridity (45), and climate change could exacerbate spring droughts by increasing the time between snowmelt and monsoonal rains (64).

### Common garden field experiments

To identify loci associated with climatic adaptation, we conducted genome-wide association studies linking genotypes to climatic niche parameters derived from large-scale multi-year common garden experiments (14). Since 2013, we have exposed multiple cohorts of seeds and seedlings from maternal lines generated from seeds collected in the Elk Mountain Range (Colorado) to manipulated snowpack and temperature treatments within the context of natural spatial and temporal variation in biotic and other abiotic conditions in five common gardens (Fig. 1 and SI Methods, ref. 14).

Anderson et al. (14) built a climate-driven integral projection model from regressions of eight vital rates measured across the life cycle and calculated the timing of snowmelt and growing season temperatures that optimized lifetime fitness (λ) for each accession. We computed eight additional parameters describing the thermal and snowmelt niche (Fig. 1; Table S17): the maximum and minimum climatic values for which λ ≥ 0.88 × the maximum lambda, sampling over variation in the other driver (65); climatic breadth (maximum - minimum value for each driver); and the two-dimensional climate niche (the area defined by the contour of the surface of λ as a function of standardized snowmelt and temperature, setting the contour equal to 0.88 × maximum λ).

### DNA extraction and sequencing

We generated low-coverage whole-genome sequence data for 219 accessions from 95 populations represented in our common garden experiment and 121 additional accessions from natural populations adjacent to each garden (Table S1 and S15; average coverage: 2.65x). DNA extraction, library preparation, read processing and alignment to the *B. stricta* SAD12v2.2 reference genome are described in SI Methods.

We performed genomic analyses in ANGSD (90), which uses genotype likelihoods rather than discrete genotype calls. Population structure was characterized using PCAngsd and the first four principal components were included as covariates in association analyses (Fig. S11).

### Genome-Environment Analysis and Genome-Wide Association Study

We conducted genome-environment association (GEA) analyses using climatic variables describing snow dynamics, temperature, precipitation, and aridity derived from regional and locally calibrated climate datasets (SI Methods; Table S18). Associations between SNPs and environmental variables were tested in ANGSD using latent genotype association models and the 95 sequenced populations. We applied the same approach to conduct genome-wide association analyses (GWAS) using the ten niche parameters for the 219 maternal lines (Table S1-S2). We used a Bonferroni correction to determine the significance threshold (P < 1.83×10^-8^). Candidate genes associated with significant SNPs were identified using a *de novo* annotation of the SAD12v2.2 reference genome constructed from gene prediction, orthology-based annotation, and functional annotation pipelines (SI Methods).

### SNPs for analyses of local adaptation and climate change responses

We identified 366 hits associated with 7 climatic niche parameters in the GWAS and 1371 hits associated with 65 climatic factors in the GEA (Table S2). To derive a set of markers to test hypotheses about local adaptation and climate change responses, we thinned these SNPs based on proximity as a proxy for linkage disequilibrium (LD) by grouping them into 10kb clusters consistent with estimates of LD decay in *B. stricta* (66) and retaining the SNP with the lowest p-value in each cluster. We excluded vapor pressure deficit or mean dew point temperature SNPs from downstream analyses as we did not measure these factors in our experiments. This process resulted in 127 GWAS SNPs and 78 GEA SNPs.

### Phenotypic effects of genotypic variation and elevational gradients in genotypic likelihoods

To determine the elevation of origin of the major allele in the thinned SNPs, we analyzed the probability of homozygosity of the major allele at each SNP as a function of source elevation or climatic factor, including a random effect for source population with a beta distribution using the R package glmmTMB ver. 1.1.13 (67). Positive slopes indicate that the major allele has a higher probability of association with accessions from high elevation populations, delayed snowmelt, or increased temperatures. For the GWAS markers, we examined variation in the mapped climate niche parameter as a function of the probability of homozygosity of the major allele, with population as a random effect using lme4 ver. 1.1.35.5 (68). We examined residuals for homoscedasticity and normality using DHARMa ver. 0.4.7 (69). We conducted univariate analyses for each SNP and applied a false-discovery rate correction (29) for multiple testing separately for the GWAS and GEA sets of markers. We then categorized markers into the most probable genotype (heterozygote or low or high elevation homozygote) based on a likelihood value of >0.7 from ANGSD, which yielded results that were concordant with analyses of the continuous probability estimates (Fig. S8).

### Genetic basis of local adaptation

We evaluated genotype × climate interactions for fitness (adult survival and seedling recruitment) to test for genetic trade-offs vs. conditional neutrality with data from the manipulative common garden experiment (N=14,170 adult transplants and 33,211 seeds with sequence and fitness data, SI Methods). To distinguish long-term adaptive divergence from short-term responses to environmental variability, we partitioned the climatic predictors into a between-environment component (mean value for each garden or garden × snow treatment combination, hereafter: mean climate) and a within-environment component (annual and microenvironmental deviations from that mean, hereafter: climate deviation) (70). This framework assessed whether local adaptation arises through long-term climatic differences across environments (significant mean climate × genotype interactions) while controlling for short-term spatio-temporal variability within environments (climate deviation).

We matched each of the 41 GWAS and 40 GEA SNPs with the climate variable for which the SNP was detected. For the GEA, we focused on five factors: the timing of snowmelt, accumulated precipitation during the growing season, mean daily temperature during the growing season, minimum daily growing season temperature (reflecting minimum night-time temperatures), and maximum daily growing season temperature (Table S19). For each SNP separately, we analyzed fitness as a function of mean climate, climate deviation, genotype, mean climate × genotype, climate deviation × genotype, cohort, the four population genetic PCs to account for population genetic structure, and random effects for garden, block nested within garden, and accession nested within population using glmmTMB ver. 1.1.13 (67) with a binomial distribution and a logit link. The random effect of garden accounted for site-level differences in fitness unrelated to climate. For annual survival, we also incorporated covariates for year of measurement and plant size in the previous year, and a random effect to control for repeated measurements of the same individual. For snowmelt timing, we modeled seedling recruitment as the probability that a seed planted in the fall of year i germinates in the spring and the seedling survives until the end of the year i+1, as germination occurs shortly after snowmelt and snow dynamics can affect seedling survival in the first year of growth. We included quadratic effects of mean snowmelt timing to account for non-linearity. For temperature and precipitation, we modeled seedling recruitment as the probability of seedling survival after germination in the spring of year i+1, as those climatic variables were averaged across the growing season.

We used a false-discovery rate correction (29) to control for multiple testing across loci within each set (GWAS vs. GEA), and pre-planned contrasts to assess the directionality of significant genotype by climate interactions (SI Methods). Open top chambers increased temperatures far beyond the bounds of historical thermal variation at these sites (14). Therefore, we excluded plants exposed to open top chambers from these local adaptation models, but we included them in our analyses of climate change responses. For models with snowmelt timing as a predictor, we retained snow removal and control (seeds and adult plants) and snow addition (seeds only) treatments. For models that included temperature and precipitation as predictors, we subset the data to include only plants exposed to ambient snowmelt.

#### Validation experiment

We validated the fitness genotype by snowmelt timing interactions from the main experiment with a reciprocal transplant experiment (14). We transplanted N=7,044 3 month old rosettes from 122 accessions into control and snow removal blocks in the five common gardens in fall 2016, monitored fitness from planting through fall of 2022 (14), monitored survival of an additional N=371 seedlings originally transplanted as seeds into the same experiment in fall 2015, and sequenced 97 accessions (19 lines per population; range 12-23, N=5,909 plants with sequence and fitness data, Tables S14-15, SI Methods). Owing to the smaller sample size and spatiotemporal sampling of climatic variables, we did not investigate genotype by temperature interactions.

### Climate change analyses

To test whether genotypes exist that could increase in fitness under the combination of projected warmer temperatures and earlier snowmelt timing, we examined fitness as a function of marker genotype by climatic interaction for all 127 GWAS and 78 GEA SNPs remaining after pruning for linkage disequilibrium. We retained the within–between decomposition (70) for the timing of snowmelt, which represents a discrete seasonal event that arises from the interaction of multiple drivers, including winter snowfall and snowpack, winter and spring temperatures, and the cumulative outcome of surface energy balance over time (e.g., 71, 72). It is inherently context-dependent: the same ordinal day of year corresponds with very early snowmelt at high elevation and very late snowmelt at low elevation. In contrast, temperature is measured on an absolute scale (°C), which has the same meaning across sites and does not require partitioning.

We analyzed fitness as a function of: temperature; partitioned snowmelt timing; SNP genotype; two-way and three-way interactions of mean snowmelt timing, temperature and genotype; two-way interactions of snowmelt deviation and genotype (to control for short-term responses to spatio-temporal variability); and all covariates (including genetic principal components) and random effects in glmmTMB (67) using binomial distributions with logit links (SI Methods). Preliminary models revealed that the quadratic effect of snowmelt timing was significantly negative after false-discovery rate correction in all but four linear models of annual survival (3 GEA and 1 GWAS SNP); we included this quadratic effect owing to higher model performance, as assessed via AICc values. There was no evidence for quadratic fitness functions for temperature. These models used data from adult plants exposed to all treatment combinations (ambient snowmelt, snow removal, ambient temperature, and elevated temperatures; N=14,170 transplants from 219 accessions), and all data from seedlings exposed to ambient snowmelt, snow removal and snow addition treatments (N=33,211 seeds from 191 accessions). Our snow removals and open top chambers reflect climate change projections for the region (14).

We estimated the fitness of each genotype at each marker under end-of-century projected snowmelt timings and temperatures in the RCP4.5 climate change scenario for each of the five experimental garden sites (SI Methods) and contrasted the predicted future fitness of the major allele homozygote against the other genotypes. We leveraged the sequence data collected from the five *B. stricta* populations adjacent to each garden to examine within-population genotypic variation at the SNPs identified in these climate change analyses (Table S14). If the allele with the predicted highest future does not currently exist in homozygous or heterozygous form within a given natural population, we classified the SNP as an <u>assisted migration locus</u> if the allele exists in either up-or downslope populations. If this allele does exist within a location but is not currently fixed there, we considered it to be an <u>evolutionary rescue locus</u> as the beneficial allele could rise to fixation as climate change proceeds.

## Supporting information

Supplemental Methods and Results

## Acknowledgments

We thank the Rocky Mountain Biological Laboratory, the Crested Butte Land Trust and the Estess family for permission for the common gardens. We are grateful to the backcountry skiers who conducted the snow removals, field researchers for their assistance collecting vital rate data, and greenhouse staff who assisted with growing plants prior to transplanting, including M. Robbins, I. Jameel, S. Day, B. Chowdhury, C. Daws, A. Battiata, C. Boerner, T. Morrison, S. Srivatsan, N. Workman, H. Nagle, M. Boyd, K. Tarner, D. Proudfoot, T. Kilgore, C. Smith, A. Beason, B. Ammon, A. Tiberio, P. Innes, K. Haner, A. Esparza, D. Eldridge, H. Grimess, E. Cheslock, E. Ross, R. MacTavish, A. Bohon, J. Adachi, J. Gall, M. Verner-Crist, M. Manto, and C. Beutler. C. Berg, H. Agha, M. Wilson Brown, and M. Wiliams commented on a previous draft. D.A.D was supported by a Plant Resilience Institute Postdoctoral Fellowship (Michigan State University). Funding came from the National Science Foundation (DEB-1553408 to J.T.A.).

## Data availability

Code and data are available on Zenodo at 10.5281/zenodo.24380391. Sequence data are available on NCBI under the BioProject accession PRJNA1564764.

## Author Contributions

J.T.A and D.A.D. designed research; J.T.A and D.A.D. performed research; J.T.A and D.A.D. analyzed data and M.L.D. calculated climate niche parameters; and J.T.A and D.A.D. wrote the paper, which M.L.D. and E.J. edited.

## Competing Interest Statement

None declared

## Notes

### Competing Interest Statement

The authors have declared no competing interest.

### Summary of Updates

Minor word changes were made throughout the manuscript to improve clarity.

