## Supplemental Methods and Results for "The genetic basis of adaptation to continuous environmental gradients informs predictions about climate change responses"

**Supporting Information Text**

**Extended methodologies**

**Common garden field experiments.** Prior to the field experiments, we collected seeds from up to 30 *B. stricta* individuals per population from sites across elevational gradients over multiple mountains in the Elk Mountain Range in Colorado (2498m-3673m; Fig.1, Table S1), and propagated these seeds in the greenhouse for at least one generation to homogenize maternal effects and generate inbred lines (hereafter: accessions). We then established fenced common gardens at five locations at different elevations near the Rocky Mountain Biological Laboratory (Gothic, CO, USA; garden elevations: 2553m, 2710m, 2890m, 3133m, 3340m). Since 2013, we have exposed multiple cohorts of seeds and seedlings from these accessions to manipulated snowpack and temperature treatments within the context of natural spatial and temporal variation in conditions to model the climatic niche integrated across the lifespan.

*Experimental manipulations I: Snowpack experiments*: We grew multiple siblings per accession for 3 months in the greenhouse before transporting juvenile rosettes to Colorado and transplanting N=6,437 individuals into the matrix of natural vegetation in control and snow removal blocks in all five gardens in October 2013 (105 accessions from 44 source populations ranging in elevation from 2694 -3690 m). Our snow removal treatment accurately captures phenological responses to climate change from longitudinal studies (1). Planting in the autumn exposes *B. stricta* individuals to overwinter vernalization cues necessary for flowering. We replicated this experiment in the fall of 2014, when we transplanted experimental N=3,403 juvenile rosettes (152 accessions from a total of 49 source populations, range of source elevations: 2499-3530 m) and N=19,151 seeds (205 accessions from a total of 56 populations, range of elevations: 2,519-3,690m) into control and snow removal treatments in all gardens. In April-June of each year from 2014 - 2022, when snowpack receded to 1 m depth in each site, we exposed these transplants to either contemporary conditions or a snow removal treatment by shoveling snow to a depth of 10 cm (2). This protocol prevents damage to plants, while hastening snowmelt, reducing soil moisture, and lengthening the dry period between snowmelt and monsoonal rains, all of which are conditions associated with and projected under climate change (3, 4).

We expanded this experiment to examine seed fitness in response to a greater range of snowmelt dynamics in fall 2016 by transplanting seeds into control, snow removal and snow addition treatments in all five gardens (N=29,676 seeds; 100 accessions from a total of 50 source populations, range of elevations: 2,499-3,530 m). Our snow addition treatment delayed snowmelt by 2-7.7 days, reflecting recent historical climates (5). In 2018, we included one new treatment, enabling us to examine seed fitness in response to control, snow removal, snow addition, and snow addition plus supplemental growing season water to simulate a benign climate in all five gardens (N=15,977 seeds; 50 accessions from a total of 50 source populations, range of elevations: 2,499 - 3623 m) (5). In this study, we exclude seeds exposed to the snow addition plus supplemental watering from fitness analyses to reduce the risk of conflating snow dynamics with precipitation.

*Experimental manipulations II:* *Multifactorial experiment of snowpack and growing season temperature*: To evaluate fitness in response to a greater range of climatic conditions, we conducted a 2 × 2 multifactorial manipulation of snowpack and growing season temperature in the falls of 2018 and 2020. In addition to snow removals on half of the experimental blocks (as described above), we established hexagonal passive open top chambers made from 55 cm tall sections of clear polycarbonate paneling, with a base diameter of 1 m and a top opening of ~55 cm (6). These open-top chambers increased average temperatures by 4.1°C over the course of this study (2). Owing to limited space in the smallest high elevation garden (3340m), we deployed pentagonal (5-sided) chambers there; with these structures, we were able to achieve similar levels of warming as the hexagonal chambers in other locations (2). We removed all chambers in the fall to prevent damage and to ensure that snow did not accumulate in these structures and replaced them on the day of the first spring census (May-June) after complete snowmelt across all blocks. We did not deploy open top chambers in our seed experiments.

In October 2018, we planted N=4,286 rosettes into all four treatment combinations of snowmelt timing and growing season temperature in all but the lowest elevation garden (N=57 accessions from 28 source populations ranging in elevation from 2499 - 3623 m, ref. , 2). In October 2020, we replicated the experiment, using all five gardens (N=4,594 rosettes; 39 accessions from a total of 38 source populations, range of elevations: 2499 - 3673 m, ref. , 2). Snowfall in the lowest elevation site was minimal during the 2020-2021 winter, and the snowbank was less than 10 cm in the spring, preventing us from doing snow removals at that site. Owing to high overwinter mortality (85.7% mortality; 788 overwinter deaths of 919 transplants) along with uneven distribution of living transplants across temperature treatments, we did not deploy warming chambers at that site. We conducted all snow and temperature manipulations at the other four gardens across the duration of the experiment.

*Data collection:* We censused the transplants 2-4 times per week from planting through 2022 to measure germination and seedling survival, plant size, survival, flowering success, and fecundity. To capture data on snowmelt timing, we directly observed the first day each experimental block was 100% snow free in the springs of 2014 and 2015; thereafter, we mounted time lapse cameras (Brinno; Taipei City, Taiwan) on fence posts and reviewed the footage to determine snowmelt timing. We interpolated air temperature data from a network of 122 weather stations and micro-meteorological sensors distributed throughout the study area (2).

In spring 2019, when we initiated the multifactorial manipulation of snowmelt timing and temperature, we recorded air temperature and relative humidity at ~25 cm above soil level every 30 minutes across the growing season with HOBO U23 Pro v2 data loggers (Onset Computer Corporation, Bourne, MA, USA) in 4-9 control blocks and 5-11 open top chamber blocks per garden (2). To avoid combining interpolated and measured air temperatures, we calculated the daily average temperature difference between control and open-top chamber blocks for each garden and each year from 2019-2022 (average of 4.55 ± 0.11 °C) (2), representing the duration of the multifactorial experiment. For each open-top chamber, we added the temperature difference based on the year and the garden for every day the chambers were deployed to determine the daily average temperature. We calculated the mean, maximum and minimum daily temperature during the growing season as an average of the daily mean, maximum or minimum temperatures from the day of first snowmelt to the day of first frost in the fall (minimum temperature falls below -1°C), which reflects *B stricta*’s annual growth and reproductive cycle. Interpolated air temperature and observed temperatures were highly correlated (β = 1.08 ± 0.013°C observed/°C interpolated, F_1,1866_**=**6528.7, p<0.0001, R^2^=0.78, N=1868 datapoints) (2). The average of the minimum daily temperatures provides insights into nocturnal temperatures, which are warming at a greater rate than diurnal temperatures (7).

**Validation dataset: Reciprocal transplant experiment.** We used our reciprocal transplant experiment (2, 5) to validate genotype by climate fitness functions at GWAS and GEA SNPs mapped for snow variables. In contrast with the main experiment, this reciprocal transplant included more accessions but fewer populations, focusing exclusively on the natural populations immediately adjacent to each common garden. After seed collection from these sites, we grew lines in the greenhouse for a generation to homogenize maternal effects and generate selfed maternal lines. We planted N=7,044 juvenile rosettes from 122 accessions (15-30 lines per source population, Table S15) into control and snow removal blocks in the five common gardens in the autumn of 2016. Gopher tunnels killed 146 of the transplants during the first winter, reducing the sample size to 6,898 individuals. We monitored fitness from planting through fall of 2022. Of the 122 total accessions transplanted into this experiment, we sequenced 95 lines (19 accessions per population; range 12-23). Thus, this validation dataset has less statistical power than the main experiment. We only analyzed fitness genotype by snowmelt timing interactions, not temperature interactions, owing to the limited sample size and shorter duration of the study. Furthermore, we have excluded the seed transplant experiment (2) from this validation effort because it only included one seed cohort planted in fall 2015. Additionally, very limited snowpack in the lowest elevation site in winter 2015 prevented us from doing snow removals there, and not a single seed germinated in that site from that experiment (2). Thus, that seed dataset included seeds exposed to a much more limited range of spatio-temporal variation in snowmelt timings than the main experiment.

**DNA extraction and sequencing**

We sequenced 219 maternal lines from the field experiment (95 populations; 2.88 ± 2.19 lines per population, mean ± standard deviation; Table S1). Given that *B. stricta* primarily self-pollinates (8), inbred full siblings share high sequence similarity (9). We extracted DNA using Qiagen DNeasy Plant Mini extraction kits (Qiagen; Hilden, Germany), and completed library preparation using the Nextera DNA Flex kits (Illumina; San Diego, CA, USA). We sequenced these accessions at the Georgia Genomics and Bioinformatics Core for 300 cycles at 150-bp using the Illumina NextSeq’s mid output paired-end sequencing protocols to obtain low coverage whole genome sequences. We also sequenced N=121 accessions from the natural populations adjacent to each garden (16-35 accessions/population, Table S15) to examine whether local populations harbor genetic variation at SNPs associated with climatic adaptation.

We trimmed Illumina Nextera adaptors and PCR contaminants from the raw reads using Trimmomatic v. 0.39 (10) with a seed mismatch count of 2, a palindrome clip threshold of 30, and a simple clip threshold of 10 (ILLUMINACLIP/NexteraPE-PE.fa:2:30:10). We performed a sliding window trimming with 4 bp windows and a minimum quality of 15. Trimmed reads were aligned to the *B. stricta* SAD12v2.2 reference genome (GenBank assembly GCA_018361395.1) (11) using BWA-MEM v. 07.17 (12). We removed duplicate reads using SAMtools v. 1.18 (13) and visualized sequence quality using Qualimap v. 2.2.1 (14).

We conducted GEA and GWAS analyses in ANGSD v. 0.940 (15), which uses a probabilistic framework to estimate genotype likelihood scores, thereby incorporating uncertainty from low sequencing depth into the analyses. We calculated genotype likelihoods with the SAMtools genotyping method (-GL 1) and the extended base alignment quality algorithm (-BAQ) (16), using a minimum minor allele frequency of 5%, with the reference genome as the known, major allele and biallelic sites with a probability of not being polymorphic at 1x10^-6^_._ Finally, sites were used if they had a minimum quality score of 20 and a minimum reference mapping score of 30. This resulted in 2,737,409 loci for use in association analyses.

To account for population structure (17), we used genotype probabilities to estimate a genetic covariance matrix between individuals in PCAngsd v. 1.2 (18), from which we derived an eigenvalue decomposition matrix with R (v.4.1.2; R Core Team, 2021). We included these top 4 PC axes in all subsequent models, as they explained 28.4% of the variation between accessions and encompassed the “elbow” of the scree (19) (Fig. S12).

**Genome-environment analysis**

Genome-environment analysis (GEA) evaluates statistical associations between environmental variables and sequence variation to detect loci that could influence climatic adaptation (17). Aridity, precipitation, snow dynamics, and growing season temperature shape fitness in *B. stricta* (2, 20–22) and other high elevation species (e.g., 23). Course-scale climate datasets such as BioClim (24) often provide less accurate precipitation estimates (25) and underestimate snowpack (26) across complex topographical regions. Therefore, we leveraged fine-grained environmental data suitable for topographically complex regions from three sources. and Furthermore, we deliberately matched accessions collected in the early 2010s (Table S1) with climate data from the same timeframe. Mismatches between the collection dates of sequenced accessions and climatic data in contemporary GEA studies could lead to the identification of SNPs associated with climatic maladaptation.

SnowClim: For each of the 95 source populations, we extracted data for 19 climatic variables from SnowClim v1.0 (27), which model snow dynamics from in the western United States, using the R package, terra v. 1.8-42 (28). We used the contemporary (CTRL) dataset to obtain annual data for total snow cover days, depth of largest snowfall events, snow duration, snow free days, minimum, mean, and maximum snow depth, minimum, mean and maximum snow water equivalent, and total snowfall over the period of 2000-2013 (Table S2 and S18). We calculated the mean value for each metric across the 13-year period to be used as phenotypes in our GEA.

PRISM: PRISM is a high-resolution gridded climate dataset that combines weather station observations with elevation-based adjustments (29) (Table S2 and S18). For each population, we extracted monthly and annual data for precipitation, minimum and maximum temperature, mean dewpoint, and minimum and maximum vapor pressure deficit data using the 30-year normals (1991-2020) (29). Additionally, we calculated means for the growing season by averaging monthly data across April through September. Collectively, the annual, monthly and growing season data resulted in 97 environmental variables for input into the GEA (Table S2 and S18).

Source climate data: The accessions transplanted into our five common gardens and sequenced for the genome-environment association study originated from un-instrumented sites in the Elk Mountains of Colorado (2). For this study, we leveraged customized calculations (presented in 2) of snowmelt timing for the source populations based on long-term Landsat 5-9 observations of normalized differential snow index (30) and interpolated air temperature estimates from a network of 122 weather stations and micro-meteorological sensors distributed throughout the study region using a Bayesian approach (31) for the current reference period of 2002-2021 at these sites. Anderson et al. (2) then defined the growing season as occurring from the estimated (posterior median) date of first snowmelt to the day of first frost in the fall (minimum temperature falls below -1°C).

**GEA and GWAS analyses**

We used environmental values from SnowClim, PRISM, and the source population data as quantitative phenotypes in ANGSD for the GEA (Table S2 and S18) and we performed GWAS with the ten climate niche parameters derived from climate-driven demographic models (Table S2 and S17). We included the 4 population structure PC variables as covariates with the 2,737,409 SNPs in a latent genotype association model (-doAsso 4), which uses a generalized linear model to estimate effect size of the genotypic probabilities given the phenotypic data and covariates (32). We analyzed each environmental variable and climate niche parameter separately. The output from ANGSD provides likelihood ratio test (LRT) values on a per-site basis. We converted these LRT values to p-values in R version 4.1.2 using the pchisq function and used a Bonferroni correction to consider hits with a p-value of 1.83x 10^-8^ or lower as significant (p=0.05/2737409 sites) for further analyses. We generated Manhattan plots using ggplot2 ver. 4.0.2 (33) and ggrastr ver. 1.0.2 (34)**.**

**Gene annotations of significant sites**

We aligned genomic data to the unannotated SAD12v2.2 *B. stricta* reference genome, developed from a Colorado accession near our lowest elevation garden (11, 35). We constructed a genome annotation by first predicting gene locations using AUGUSTUS v3.5 (36) and OrthoFinder v3.1 (37) to identify orthologous genes from seven Brassicales: *Arabidopsis thaliania*, *Brassica oleracea*, *Cardamine amara*, *Capsella rubella*, *Eutrema salsugineum, Microthlaspi erracticum* and *Thalspi arvense* (Table S20). We supplemented our annotation with BLAST+ v2.16 (38), InterProScan v5.74 (39), and eggNOG-mapper v2.12 (40). Our assessment using BUSCO v5.8.3 (41) found 96.1% completeness when compared to the Brassicales_odb10 dataset.

**Phenotypic effects of genotypic variation and elevational gradients in genotypic likelihoods**

The GEA identified 1165 SNPs associated with macroclimatic variation and the GWAS identified 366 SNPs associated with the thermal and snowmelt niche. We then pruned these hits for linkage disequilibrium, and examined elevational, climatic and trait clines for 127 SNPs detected via GWAS and 78 SNPs detected via GEA. As the probability of major allele homozygosity is a proportion on the [0,1] scale, we first applied a beta transformation (42) to this response variable, which we then analyzed as a function of source elevation or climatic factor, including a random effect for source population with a beta_family() distribution in the glmmTMB function of R glmmTMB ver. 1.1.13 (43).

*SNPs identified via GWAS*: In 162 separate tests, we found 105 significant relationships after false-discovery rate correction for multiple testing (44) at 85 unique SNPs for the relationship between the probability of homozygosity of the major allele and variation in one or more climate niche parameters (Table S5). Additionally, 41 SNPs showed significant elevational clines in the probability of homozygosity of the major allele such that we could define alleles as coming from either low or high elevation source populations (Fig. S6; Table S4). These elevational patterns were all concordant with clines in climatic niche parameters and climatic gradients in this geographic region, and thus consistent with expectations under local adaptation (e.g., Figs.1B and S6).

*SNPs identified via Genome-Environment Association (GEA)*: From 421 analyses of SNP/climate combinations, we detected 243 significant relationships between allelic variation at 57 unique loci and climatic factors after correction for multiple testing (Table S6), and 40 SNPs exhibited significant associations between the probability of homozygosity of the major allele and source elevation (Table S7). We found significant clines in one or more climatic variables for all 40 SNPs displaying elevational clines, and the direction of the elevational gradient was concordant with the direction of the climatic gradient (e.g., Fig. S6, and Table S6 and S7). Thus, across both analyses, elevational variation in genotypic probabilities was concordant with trait and climatic variation, such that low elevation alleles were associated with early snowmelt timing, warmer temperatures, and decreased precipitation.

**Genetic basis of local adaptation.**

We matched each of the 41 GWAS and 40 GEA SNPs with the climate variable for which the SNP was detected (GEA) or for the climate variable associated with the climate niche parameter (temperature or snowmelt timing in the GWAS). Overall, the GEA SNPs were associated with 37 climatic factors, and many loci showed significant associations with multiple correlated variables. For instance, SNP 1_21062864, was significantly associated with 11 maximum temperature variables (e.g., maximum temperature in March, April, May, June, July, August, September, October, November, as well as the average maximum temperature annually and during the growing season), 9 mean temperature variables, 2 snowmelt variables and 2 minimum temperature variables. To avoid redundant analyses of fitness genotype by climate interactions, we condensed these variables to five climatic factors (Table S19): (1) the timing of snowmelt, (2) accumulated precipitation during the growing season, (3) mean daily temperature during the growing season, (4) minimum daily temperature averaged across the growing season (reflecting minimum night-time temperatures), and (5) maximum daily growing season temperature. We excluded plants in the open top chambers from local adaptation models because the chambers increased temperatures far beyond the bounds of historical thermal variation in these sites (2).

After partitioning observations of each climatic predictor into a between-environment component (climate mean) and a within-environment component (climate deviation), we constructed separate for loops in R with the set of GEA and the set of GWAS SNPs, looping over all SNPs separately to conduct generalized linear mixed models that analyzed fitness as a function of partitioned climatic variation (mean climate and deviation from the mean), SNP genotype, their two-way interactions (mean climate × SNP genotype and climate deviation × SNP genotype), planting cohort, year of measurement, the four population genetic PCs generated from ANGSD to account for population genetic structure. These models included random effects for garden, block nested within garden, and accession nested within population using the glmmTMB function of the R package glmmTMB ver. 1.1.13 (43) with a binomial distribution and a logit link. We standardized continuous fixed effects to a mean of 0 and standard deviation of 1, but we built figures on the raw scale using ggeffects ver. 2.3.2 (45) and ggplot2 ver. 4.0.2 (33). For annual survival, we also incorporated a covariate for plant size in the previous year, and random effects for unique plant identifier and year to account for repeated measurements on the same individual. Seedling recruitment is defined as the probability that a seed planted in the fall of year i germinates in the spring of year i+1 and the seedling survives until the end of the year i+1. Seeds that fail to germinate could remain dormant, although dormancy is low in this system (5, 46). For models of seedling recruitment, we included quadratic effects of mean snowmelt timing to account for non-linear fitness functions. We used a false-discovery rate correction (44)to account for multiple testing across loci within each set of markers (GWAS vs. GEA).

When convergence issues occurred under the default optimizer, we refit models using an alternative optimizer (BFGS); all reported results are from converged models with positive-definite Hessians. For 2 GEA SNPs (3_14426161 and 4_18823142) for which we modeled seedling recruitment as a function of minimum temperature, convergence problems persisted even with the alternative optimizer; in those two cases, we achieved model convergence by replacing the random effect of genotype nested within population with a simpler random effect of genotype. For 10 GWAS SNPs and 15 GEA SNPs, only 1-2 maternal lines were heterozygous or carried the minor allele homozygote. In those cases, we combined heterozygotes and minor allele homozygotes into one category for comparison against major allele homozygotes.

*Contrasts for assessing directionality of genotype by climate interactions:* We used a series of pre-planned contrasts to assess the directionality of genotype by climate interactions, and we extracted the slopes of the relationship between climatic variables and genotypes using the emmeans package ver. 1.10.3 (47) to detect genetic variation in the fitness consequences of climate change. To assess local adaptation, we first contrasted the relative odds (and 95% confidence interval) of adult survival or seedling recruitment for the low elevation homozygote vs. the high elevation homozygote at the earliest mean snowmelt timing, the hottest mean temperature or the lowest mean accumulated precipitation in the dataset. Similarly, we calculated the relative odds of survival and recruitment for the high elevation homozygote vs. the low elevation homozygote at the most delayed mean snowmelt timing, lowest temperature, and greatest accumulated precipitation. At certain SNPs, no accession carries the minor allele homozygote; in those cases, we constructed our models to contrast the major allele homozygote vs. the heterozygote.

If local adaptation is driven primarily by genetic trade-offs, the odds ratios would significant exceed 1 for both of our contrasts, such that the low elevation allele outperforms the high elevation allele in hot and arid climates associated with low elevation environments, and high elevation alleles would outperform low elevation alleles in cool and mesic climates. In the case of conditional neutrality, either low or high elevation genotypes would have elevated fitness in the respective home climate without experiencing a fitness cost in the opposing environment. In our models, that would manifest as the odds ratio significantly exceeding 1 for one of the contrasts while the other contrast would not show a statistical difference in the fitness of the low and high elevation alleles. If both contrasts are significantly lower than 1, we would infer local maladaptation (i.e., the high elevation allele had greatest performance in low elevation climates and vice versa). Finally, a significant genotype by mean climate interaction could arise if one genotype outperforms the other in all climates but has a more pronounced advantage in either high or low mean climate. In our models, this pattern would result in a relative odds ratio >1 at one end of the climatic spectrum and <1 at the other end.

For all models of seedling survival, we recovered negative quadratic effects of mean snowmelt timing, and the vertex (=optimum) occurred within the range of snowmelt timings, suggesting that intermediate snowmelt times optimized fitness. In that case, it is no longer possible to distinguish between conditional neutrality and genetic trade-offs using the approach described above, as the relative fitness of genotypes depends on the exact climatic value being considered. Therefore, we computed the optimal timing of snowmelt (and 95% confidence interval) for each genotype as the climatic condition at which the negative quadratic fitness function peaked. We expected low elevation genotypes to have optimal fitness at earlier snowmelt timings and higher temperatures than high elevation genotypes.

We also extracted the slopes of the relationship between climatic variables and fitness for each genotype at each SNP using the emtrends function of using the emtrends function of the emmeans package ver. 1.10.3 (47) to examine how the odds of survival and seedling recruitment vary across the observed climatic gradients. For each model for which we detected a significant SNP by climate interaction after Benjamini-Hochberg (44) correction for multiple testing, we determined whether fitness increased (odds ratio > 1) or decreased (odds ratio <1) with the climatic factor for each genotype to evaluate predicted fitness in future climate scenarios (early snowmelt, increased temperatures, reduced precipitation) and to ascertain whether the fitness consequences of climate change differ for loci displaying local adaptation vs. conditional neutrality.

*Validation dataset*: For the reciprocal transplant experiment, we analyzed adult survival as a function of mean snowmelt timing, deviation from the mean snowmelt timing, SNP genotype, two-way interactions of SNP genotype and mean snowmelt timing and SNP genotype and deviation from the mean, including covariates for year, plant size in the previous year, and random effects for genotype nested within source population, garden, block and plant identifier. We focused on the 24 GWAS SNPs associated with the snowmelt niche and the 2 GEA SNPs associated with snow-related climatic factors. Given that this experiment only included accessions originating from each of the five common gardens, we excluded genetic PC variables from this model.

**Climate change models**

We aimed to detect genotypic variation at the GEA and GWAS SNPs that could enhance fitness under climate change scenarios. Unlike the local adaptation analysis, these models did not require information on the elevational origin of the major allele; therefore, we leveraged the full suite of markers that we identified after pruning for linkage disequilibrium. A single SNP (1_5794422) was recovered in both GWAS and GEA analyses; we have included this SNP with the GWAS set of markers. Furthermore, we excluded 1 GEA SNP (1_13477980) from analysis of seedling recruitment as it had limited genotypic variation (only three accessions were homozygous for the major allele with the remaining accessions being heterozygous).

We constructed for loops in R to model annual adult survival and juvenile recruitment as a function of temperature, partitioned snowmelt timing, the categorical genotype at each SNP, climatic factor by genotypic interactions, and a three-way interaction of mean snowmelt timing × temperature × SNP genotype, with covariates for the four genetic principal components, plant size in the previous year (survival models only), cohort, and random effects for accession nested within source population, experimental block, garden, year (survival models only) and unique plant identifier (survival models only) in glmmTMB (43) using binomial distributions with a logit link. The three way interaction of temperature, mean snowmelt timing, and marker genotype enabled us to predict future fitness for all genotypes at all SNPs in response to multivariable climate change, as temperatures are increasing and snowmelt timing is advancing simultaneously. We incorporated quadratic effects of mean snowmelt timing into all but four models to account for non-linearity in fitness responses to this climatic factor. Finally, the inclusion of deviation from mean snowmelt timing and the two way interaction of this factor and SNP genotype enabled us to control for fitness responses to short-term temporal or spatial variability in conditions.

We reverted to an alternative optimizer (BFGS) if models with the default optimizer failed to converge (1 GWAS and 1 GEA SNP for survival). For 33 SNPs detected via GWAS and 28 SNPs detected via GEA, very few accessions in our dataset were homozygous for the minor allele. For these SNPs, we categorized allelic variation as: homozygous for the major allele vs. at least one minor allele (including heterozygotes and accessions homozygous for the minor allele).

We used a likelihood ratio test to contrast our full model for each SNP against a null model that excluded the interactions of SNP genotype with mean snowmelt timing and with temperature. This omnibus test evaluates the significance of the interaction of genotype with either or both climate variables for each SNP. We applied a Benjamini-Hochberg (44) correction for multiple testing across the GEA and GWAS SNPs combinations to this omnibus p-value. For markers that demonstrated significant genotype by climate interactions through this omnibus test, we extracted the slope of the relationship (the odds ratio) between fitness and climate for each available genotype (major allele homozygote, heterozygote and minor allele homozygote) using the emtrends function of the emmeans package ver. 1.10.3 (47) to evaluate how the odds of adult and seedling survival vary across the observed climatic gradients for each genotype. Odds ratios (OR and 95% CI) > 1 indicated enhanced odds of survival/recruitment under projected future elevated temperatures, whereas OR < 1 indicated greater fitness under early snowmelt and reduced precipitation, reflecting climate change projections (Table S12). For all models in which we included a quadratic effect of mean snowmelt timing, we recovered negative quadratic relationships, enabling us to extract the optimal snowmelt timing (the climate under which fitness peaked) and 95% confidence interval for each genotype at each SNP (Table S13).

*Assisted migration vs. evolutionary rescue loci*: These analyses detected whether the regional pool of accessions in this study include alleles that could enhance survival under climate change; however, adaptation occurs at the population-level. To determine whether local populations harbor potentially adaptive genetic variation that could reduce the risk of extinction in climate change scenarios, we leveraged the sequence data collected from the five local *B. stricta* populations native to each garden location to examine the extent of genotypic variation within each population at the SNPs identified in these climate change analyses (Table S14).

To identify loci that could contribute to evolutionary rescue or assisted migration, we calculated the predicted fitness of all genotypes at all SNPs under future climates projected using the RCP4.5 end-of-century climatic change scenario (Table S14) (2) for each of the five experimental garden locations, using the emmeans function of the emmeans package ver. 1.10.3 (47). For each SNP displaying two- or three-way interactions of SNP marker genotype with mean snowmelt timing and temperature, we contrasted the fitness of the major allele homozygote against the other genotypes using a Benjamini-Hochberg (44) correction for multiple testing to identify the genotype with the highest predicted fitness under this projected climate scenario at the site of each experimental garden. This process identified 57 SNPs (47 from the analysis of annual survival using SNPs discovered via GWAS and 10 from the analysis of seedling recruitment using GEA SNPs).

We then examined genotypic frequencies at these 57 SNPs in the five natural *B. stricta* populations adjacent to each common garden to determine whether the genotype with the highest predicted future fitness currently exists in those populations (Table S18). For 10 SNPs, there was no genetic variation within the pool of accessions from the five local populations. We classified the remaining SNPs as assisted migration and/or evolutionary rescue loci. If the allele with a future fitness advantage does not currently exist within a local population but occurs in at least one of the other populations, we deemed that SNP to be an *assisted migration locus*, as translocation from lower or higher elevation populations could introduce this allele into the local population. In contrast, if the allele with a future fitness advantage currently exists at low frequency or exclusively in heterozygous form, we deemed that SNP to be an *evolutionary rescue locus*.

**Supplemental Results**

**Genetic basis of local adaptation**

Genotype by climate interactions significantly influenced fitness in 85 of 210 total analyses (Table S8). The number of analyses exceeded the total number of SNPs × 2 fitness components, as 3 SNPs from the GWAS mapped to both snowmelt timing and mean growing season temperature, and 15 SNPs from the GEA mapped to 2-4 climatic factors; the fitness effects of genotypic variation at these 18 SNPs were, therefore, modeled in 2-4 single climate analyses. For the SNPs identified via GWAS, significant genotype by climate interactions for fitness arose primarily in response to variation in snowmelt timing (19 SNPs) and less so to mean growing season temperature (6 SNPs). In contrast, for the GEA SNPs, we detected significant genotype interactions for four climate variables (2 SNPs for snowmelt timing, 3 SNPs for maximum temperature, 23 SNPs for minimum temperature, and 15 SNPs mean temperature), but not for accumulated precipitation (Table S8; Fig 3). This breakdown is consistent with the large number of SNPs identified for minimum temperature variables (Table S2).

Conditional neutrality can only underlie local adaptation if different loci show opposing patterns (48, 49), which we find within our dataset. For example, the high elevation homozygote is conditionally beneficial under delayed snowmelt and cool temperatures for SNP 1_14098322 whereas the low elevation homozygote at SNP 1_6920936 is conditionally beneficial under early snowmelt and warmer temperatures (Table S8).

*Local maladaptation:* We found 17 unique SNPs with local or conditional maladaptation identified through the GEA. Of these, 10 SNPs were intragenic and 7 were intergenic. Eight SNPs were either unannotated or have unknown functions, while the remaining are associated with gene repair and protein turnover (Table S16).

*Validation dataset:* With the reciprocal transplant experiment, we tested for local adaptation in the 24 GWAS and 2 GEA SNPs associated with snow variables. After false-discovery rate correction (44), we identified six SNPs with significant snowmelt timing by genotype interactions for adult survival and six additional SNPs with marginally significant interactions (FDR corrected p-value <0.08). The significant SNP from the GEA (2_16311506) showed adaptive conditional neutrality, with the low elevation homozygote outperforming the heterozygote at early snowmelt and no difference in survival at delayed snowmelt (mean snowmelt × marker genotype, χ²=6.52, p=0.0106, FDR-corrected p=0.049, Table S9). We also found a significant interaction of snowmelt timing × genotype for SNP 2_16311506 genotype for annual survival in the main experiment (χ²=12.26, FDR-corrected p=0.00404, Table S8), indicative of adaptive conditional neutrality in the opposite direction, with the heterozygote outperforming the low elevation homozygote under delayed snowmelt timing (Fig. S10).

Of the 5 GWAS SNP that displayed significant snowmelt timing by genotype interactions (Table S9), three showed adaptive conditional neutrality, consistent with the pattern from the main dataset for SNP 4_8495908 and in the opposite direction as the main dataset for SNP 1_17015801. The third SNP (7_23849428) exhibiting conditional neutrality in the validation dataset (mean snowmelt × marker genotype, χ²=13.3, p=0.0013, FDR-corrected p=0.034, Table S9) showed only marginally significant genotype by snowmelt interactions in the main dataset after correction for multiple testing (χ²=6.32, p=0.0425, FDR-corrected p=0.078, Table S8). However, the directionality of conditional neutrality reversed across datasets, with the low elevation allele being conditionally advantageous in the validation experiment and the high elevation allele being marginally conditionally beneficial in the main experiment. For the remaining two GWAS SNPs with significant genotype by climate interactions, one (6_12989886) showed apparent maladaptation and the other (3_16974565) had crossing reaction norms, but no clear conditional neutrality or genetic trade-offs. For both of these loci, the confidence intervals of contrasts between the low and high elevation homozygotes at opposite ends of the climatic space spanned one, suggesting weak overall patterns in extreme climates.

Given the limited statistical power of the validation experiment, we elected to examine the 6 GWAS SNPs with marginally significant snowmelt timing by genotype interactions (FDR-corrected p-value <0.08, all raw p-value <0.04; Table S9). In the main experiment, we found genetic trade-offs for annual survival for one (1_10026491) of these SNPs and conditional neutrality for the remaining 5 SNPs (Table S8). In the validation dataset, four SNPs have conditional neutrality, one appears as if it may be subject to genetic trade-offs (1_6920936), and the final SNP had more nuanced patterns with no clear evidence for conditional neutrality or genetic trade-offs.

**Climate change analyses**

Amongst the suite of 127 GWAS and 78 GEA SNPs retained after subsetting for linkage disequilibrium and polymorphism, 134 SNPs showed a significant main effect of snowmelt timing or temperature, and an additional 36 SNPs showed evidence of genotype by climate interactions for fitness after FDR correction (Tables S11-S13). At the SNPs displaying genotype by climate interactions for fitness, we identified 43 SNPs for which the major allele homozygote is predicted to have significantly higher fitness than the other genotypes in projected future climates primarily in the two highest elevation gardens, 3 SNPs for which the heterozygote or minor allele homozygote is predicted to outperform the other genotypes under projected climates, and 1 for which the major allele homozygote is predicted to be the best genotype in one site and the minor allele homozygote or heterozygote is predicted to be the best genotype in another site. Of these 47 total SNPs, 4 were detected via GEA and 43 via GWAS. We only found evidence for genotypic variation in projected future fitness in the three lower elevation gardens at 9 SNPs, which is consistent with worsening conditions in low elevation sites genome-wide (2).

**Figures**

**Fig. S1.** Predictions for genotypic responses to climatic variation under (A, B, D, E) conditional neutrality and (C, F) genetic trade-offs. In this system, temperatures decline with elevation and the timing of snowmelt becomes more delayed; thus, we expect the fitness of low elevation homozygotes, which evolved under warmer temperatures with early snowmelt, to increase with increasing temperatures and diminish with delayed snowmelt timing. Conversely, we predict that the fitness of high elevation homozygotes would decline with increasing temperatures and increase with delayed snowmelt. Conditional neutrality occurs if the local allele has a fitness advantage in its home site, but there is no difference in fitness of the alleles in the alternative location. In contrast, under genetic trade-offs, the local allele would maintain a fitness advantage in each environment, while also displaying higher fitness in the home vs. away site, consistent with both the local-foreign and home-away criteria of local adaptation (50). Red arrows indicate the necessary statistically significant differences in fitness of low vs. high elevation homozygotes to render a decision of conditional neutrality vs. genetic trade-offs.

**Fig. S2.** Climate varies predictably across elevational gradients, with hot low elevation sites experiencing earlier snowmelt and cool high elevation sites experiencing delayed snowmelt. Plotted are fine-grained snowmelt and temperature variables calculated for the source populations used in the common garden experiments based on long-term Landsat 5-9 observations of normalized differential snow index and interpolated air temperature estimates from a network of 122 weather stations and micro-meteorological sensors distributed throughout the study region (30), with the five common gardens displayed with red datapoints. (A) The range of snowmelt timings (the latest ordinal day of year of snowmelt - the earliest ordinal day of year of snowmelt at a given site). The (B) average (C) latest and (D) earliest ordinal day of snowmelt all increase with source elevation. Although the (E) range of temperatures (hottest - coolest temperature) increases with elevation, (F) the maximum, (G) mean and (H) minimum temperatures in the growing season all decline with elevation.

**Fig. S3. SnowClim climatic variables show clinal variation across elevation, with higher-elevation sites receiving more snow than lower-elevation sites.** Values for each of the 95 source populations were extracted from the SnowClim v1.0 contemporary (CTRL) dataset (27), which covers 2000–2013, and plotted against elevation. (A–C) Maximum, mean and minimum snow water equivalent (m). (D–F) Maximum, mean and minimum snow depth (m). (G) Mean monthly total snowfall (m). (H) Depth of the largest snowfall event (m). (I) Mean monthly number of days with snow depth >10 mm. (J) Snow duration, the number of days between the start and end of snow cover. (K) Snow-free days, the number of days without snowpack between the start and end of snow cover. Each point represents one population, and the five common gardens are shown in red.

**Fig. S4.** We extracted 98 climatic variables from PRISM, a high-resolution gridded climate dataset which combines weather station observations with elevation-based adjustments (29). This figure demonstrates elevational variation in mean growing season (April - September) PRISM climate data using the 30-year normals (1991-2020) for: (A) accumulated precipitation, (B) dew point temperature, (C) maximum temperature, (D) mean temperature, (E) minimum temperature, (F) maximum Vapor Pressure Deficit, and (G) minimum Vapor Pressure Deficit. The five common gardens are displayed with red datapoints.

**Fig. S5Manhattan**.

Manhattan plots for all GEA analyses using PRISM, SnowClim, and source climate variables, as well as GWAS climatic niche variables. Significance is displayed on the -log10 scale. Dotted lines represent a significance threshold based on Bonferroni correction (-log10(p) = 7.738).

**Fig. S6**. Elevational, climatic, and trait clines in genotypes at a representative SNP (1_6920936) discovered in the genome-wide association of climate niche parameters. (A) The probability of homozygosity of the major allele declines at this SNP declines with source elevation, consistent with (B) categorization of genotypic variation, indicating that the major allele is associated with low elevation environments. Congruent with climatic gradients in this region, the major allele has an increased likelihood of occurring in areas with (C and D) earlier snowmelt and (E and F) warmer temperatures. We found consistent clines in niche parameters, such that the major allele homozygote is associated with increased (G and H) optimal temperatures and (I and J) earlier snowmelt timings. Major = Major allele homozygote; Het = Heterozygote; Minor = Minor allele homozygote.

**Fig. S7**: Conditional neutrality at 8 GWAS SNPs demonstrating statistically significant genotype by climate interactions for annual adult survival after false-discovery rate correction for multiple testing. Ascertainment bias may restrict the determination of genetic tradeoffs because the local genotype needs to display significantly enhanced fitness in its home site relative to the foreign genotype in at least two environments (in our case: at opposite ends of the climatic gradients). The high elevation homozygote (or heterozygote) had a significant fitness advantage over the low elevation homozygote at delayed snowmelt timing for these SNPs: (A) 1_7722280, (B) 1_7848217, (C) 2_18963993, and (D) 3_16974565, but we were unable to detect a statistically significant difference between low and high elevation genotypes at early snowmelt at these loci. We detected significant conditional neutrality at the cooler end of the temperature gradient for (E) SNP 1_14098322, and at the warmer end of the spectrum for (F) 1_6920936, (G) 5_17217251, and (H) 7_9449114. Data points and regression lines are colored by genotype. To depict genotypic variation in fitness functions for adult survival, we have truncated the y-axis to 0.5 - 1, with data points for dead plants shown at y=0.5 instead of y=1, but all analyses were done with generalized linear mixed models with binomial distributions at logit links with raw data (y=0 for dead and 1 for surviving plants). Red arrows indicate at which end of the climatic spectrum we recovered a significant advantage of the local allele.

**Fig. S8**: Conditional neutrality at 6 GEA SNPs, illustrating potential ascertainment bias against the discovery of genetic trade-offs. The heterozygote had significantly higher survival than the low elevation homozygote at delayed snowmelt timing for (A) 1_21062864 and (B) 2_16311506, but we found no fitness differences at early snowmelt. In contrast, the low elevation homozygote outsurvived the heterozygote at the warmer end of the temperature gradient for (C) 2_18555145 and (D) 3_16655604, but the fitness difference was non-significant at the colder end of the gradient. Finally, we found that the heterozygote seedlings had greater establishment success than the lower elevation homozygote at cold temperatures for (E) 3_26270349 and (F) 4_13558274, with no evidence for seedling performance differences the warmest temperatures. Data points and regression lines are colored by genotype. To depict genotypic variation in fitness functions for adult survival, we have truncated the y-axis to 0.5 - 1, with data points for dead plants shown at y=0.5 instead of y=1, but all analyses were done with generalized linear mixed models with binomial distributions at logit links with raw data (y=0 for dead and 1 for surviving plants).

**Fig. S9:** Patterns of conditional neutrality differed in directionality between the provenance trial experiment (left-hand column) and the validation reciprocal transplant experiment for these four SNPs (A and B) 1_17015801, (C and D) 2_16311506, (E and F) 2_13880462, (G and H) 3_8350771. Red arrows indicate the statistically significant difference between the low elevation homozygote and the heterozygote. Survival was higher in the provenance trial experiment; therefore, we truncated the y-axis to [0.5,1] to display the patterns more clearly.

**Fig. S10:** For loci with significant snowmelt timing by genotype interactions, reaction norms typically cross at intermediate climates, reflecting historical conditions at mid-elevation sites, as exemplified by the following SNPs (A) 1_10026491; (B) 1_21062864; (C) 1_7483312; (D) 4_11395299; (E) 6_13011701; (F) 7_10270795. The X axis displays current (circle) and end-of-century RCP 4.5 projected future (arrow) snowmelt timings at the mid-elevation common garden site (pink; 2890 m) and a lower elevation garden site (orange; 2710 m). Two SNPs are noteworthy. At (D) 4_11395299, the heterozygote maintains the highest fitness across the full range of climatic variation; however, our sequencing efforts in the local populations adjacent to each garden (Table S14) recovered only one heterozygote at the highest elevation location, with the low elevation homozygote fixed in the lower four populations and the high elevation homozygote having the greatest genotypic frequency at high elevations. At (E) 6_13011701, the low elevation homozygote has the lowest fitness across the full range of snowmelt timings, but the heterozygote has a fitness advantage under projected future climates at low elevation sites.

**Fig. S11.** Principal component analysis used to account for population structure. (A) Projection of accessions onto the first two principal components derived from the genetic covariance matrix estimated from genotype probabilities in PCAngsd. Each point represents a single accession. (B) Scree plot showing the eigenvalues associated with the first 10 principal components. The first four PCs encompassed the elbow of the scree plot and together explained 28.4% of the genetic variation among accessions; these PCs were included as covariates in all subsequent analyses.

**Tables**

**Table S1: Accessions included in the genome-wide association study (GWAS) of climate niche parameters.** We provide the unique identifier for each population and accession, along with the latitude and longitude of the population, the elevation of the population, and the year in which seeds were collected from that population. In some cases, we do not have the exact year of collection, but we know a range of years within which the collection was made.

**Table S2: SNPs significantly associated with climate niche parameters (GWAS) or climatic factors (GEA).**

**Table S3: Top candidate gene for all SNPs identified in the GWAS and GEA.** Distance is distance to gene and Blast description is the description for the top gene identified through BLAST search

**Table S4: Climate niche variation at SNPs detected in the genome wide association study.** We evaluated the relevant climatic niche parameter as a function of the probability of homozygosity of the major allele in linear mixed models. This table displays each SNP, the climatic niche parameter used, the relevant statistics (Chi-squared, raw p-value, and false-discovery corrected p-value for 162 separate tests) along with the slope of the relationship between the probability of major allele homozygosity and climate and the standard error of of this slope. We note that all climate niche parameters were derived from climate-driven demographic models (integral projection models) that evaluated fitness as a function of snowmelt timing and temperature in the large-scale provenance trial experiment.

**Table S5: The probability of homozygosity of the major allele varied as a function of source elevation for 41 of the 127 SNPs identified via GWAS in a beta regression framework.** We provide the coefficient (slope of the relationship between elevation the this probability, expressed in log-odds), the standard error of the coefficient, the statistics (Chi-squared value, raw p-value and the p-value after false discovery rate correction, Benjamini-Hochberg, 1995) for separate analyses of these 83 SNPs. If the slope of the relationship is negative, then the probability of homozygosity of the major allele is greatest at low elevation sites and we define the major allele as being associated with low elevations. We conducted local adaptation analyses of fitness using the 57 SNPs for which we could define a clear elevational association with the major allele (in bold typeface).

**Table S6: Climate variation at SNPs detected in the genome-environment association**. We evaluated the relevant climatic factor as a function of the probability of homozygosity of the major allele in linear mixed models. This table displays each SNP, the climatic variable used and the major climatic group to which the variable belongs, the relevant statistics (Chi-squared, raw p-value, and false-discovery corrected p-value for 286 separate tests) along wtih the slope of the relationship between the probability of major allele homozygosity and climate and the standard error of this slope.

**Table S7: The probability of homozygosity of the major allele varied as a function of source elevation for 40 of the 78 SNPs identified via GEA in a beta regression framework.** We provide the coefficient (slope of the relationship between elevation the this probability, expressed in log-odds), the standard error of the coefficient, the raw p-value and the p-value after false discovery rate correction (Benjamini-Hochberg, 1995) for separate analyses of these 78 SNPs. If the slope of the relationship is negative, then the probability of homozygosity of the major allele is greatest at low elevation sites and we define the major allele as being associated with low elevations. We conducted local adaptation analyses of fitness using the 40 SNPs for which we could define a clear elevational association with the major allele (in bold typeface).

**Table S8: Evaluation of genotype by climate interactions for 41 SNPs detected via GWAS and 40 SNPs identified through the GEA.** This subset of SNPs showed strong evidence for elevational clines, such that we could easily identify whether the probability of homozygosity for the major allele was greater for high vs. low elevation accessions. A significant interaction between marker genotype and mean climatic factor value indicates that genotypes differ in their fitness responses to climate. We used a false-discovery rate (Benjamini and Hochberg, 1995) correction for multiple testing across markers from the same fitness component and SNP panel (GWAS vs. GEA) to assess significance of that interaction. To disentangle local adaptation from conditional neutrality and other patterns, we contrasted the odds of survival or seedling recruitment for the low elevation genotype vs. the high elevation genotype at the earliest snowmelt, warmest temperature, or lowest precipitation. We then contrasted the odds of survival or seedling recruitment for the high elevation genotype vs. the low elevation genotype at the most delayed snowmelt, coolest temperature, or highest precipitation. Odds ratios > 1 indicate that the local genotype outperforms the foreign genotype in the home climate. The final column provides our interpretation of the evolutionary pattern at each marker.

**Table S9: Evaluation of genotype by climate interactions for 24 SNPs detected via GWAS and 2 SNPs identified through the GEA for snowmelt timing in the validation reciprocal transplant experiment.** This subset of SNPs showed strong evidence for elevational clines, such that we could easily identify whether the probability of homozygosity for the major allele was greater for high vs. low elevation accessions. A significant interaction between marker genotype and mean climatic factor value indicates that genotypes differ in their fitness responses to climate. We used a false-discovery rate (Benjamini and Hochberg, 1995) correction for multiple testing across all markers (GWAS and GEA) to assess significance of that interaction. To disentangle local adaptation from conditional neutrality and other patterns, we contrasted the odds of survival or seedling recruitment for the low elevation genotype vs. the high elevation genotype at the earliest snowmelt, warmest temperature, or lowest precipitation. We then contrasted the odds of survival or seedling recruitment for the high elevation genotype vs. the low elevation genotype at the most delayed snowmelt, coolest temperature, or highest precipitation. Odds ratios > 1 indicate that the local genotype outperforms the foreign genotype in the home climate. The final column examines whether patterns were consistent with the main dataset. Given that we only had genomic and fitness data from 95 accessions in this reciprocal transplant experiment, we have only examined adult survival as a function of snowmelt timing by marker genotype interactions. We highlight statistically significant markers after FDR correction in yellow and those markers that are marginally significant (FDR-corrected p<0.08) in pink.

**Table S10: Candidate genes for SNPs displaying local adaptation (genetic tradeoffs/antagonistic pleiotropy or conditional neutrality)**

**Table S11: Results of models examining climate change responses at 127 GWAS and 78 GEA SNPs using data from all treatments, including open top chambers.**

**Table S12: Slope of the relationships between fitness (response variable) and mean temperature or snowmelt timing (for linear models only) for each genotype of the SNPs detected via GWAS or GEA.** We deemed the slope to be significant if the false-discovery rate (FDR) p-value was significant for both the omnibus test and the interaction of climatic factor(s) and marker genotype. We then assessed whether climate change could increase or decrease fitness based on the direction of the slope (outcome and climate change implication).

**Table S13: Optimal snowmelt timings extracted from climate change models.** All but four SNPs showed a non-linear relationship between snowmelt timing and fitness, such that we included a quadratic effect of mean snowmelt timing into our models. All models showed significant negative quadratic effects, enabling us to extract the optimal snowmelt timing and 95% confidence interval, which reflect the timing of snowmelt with the highest fitness in our dataset.

**Table S14: Sequence variation in the five natural populations adjacent to each common garden along with climatic conditions and projections from those sites.** Genotypic counts (not frequencies) within local populations at SNPs associated with climate niche parameters in the GWAS and climatic factors in the GEA, along with an indication of whether genetic variation exists within populations (is more than one genotype present) and if so, whether the heterozygote is present or not. The table is sorted by SNP identifier. The table indicates whether our fitness models uncovered marker genotype by climate interactions for each SNP (Genotype x Climate interactions), which genotype is predicted to have a fitness advantage over the other genotypes in end-of-century temperatures and snowmelt timings projected via the RCP4.5 climate change scenario (Fitness pattern), and whether a marker in a given location could be described as an assisted migration locus (the best genotype in future climates does not yet occur in that site) or evolutionary rescue locus (the best genotype in future climates exists in that site but is not yet fixed; Assisted migration or evolutionary rescue locus?). The smaller inset table indicates the number of maternal lines sequenced per local population and the number of maternal lines included in the reciprocal transplant experiment (the validation experiment).

**Table S15: Accessions sequenced from natural populations adjacent to each common garden**: (1) to assess whether standing genetic variation could enhance fitness in future projected climates and (2) to validate snowmelt timing results from the main genome-wide association study (GWAS) by linking genotype to fitness patterns from a reciprocal transplant experiment. We were unable to include all sequenced accessions in the reciprocal transplant experiment. This sheet indicates whether a sequenced accession was also in that experiment. In addition, some accessions in the reciprocal transplant experiments were not sequenced. The inset table shows the current (2014-2022) and end-of-century projected snowmelt timings and temperatures under a moderate climate change scenario (RCP 4.5 or Representative Concentration Pathway 4.5) based on measurements (current) and projections described in Anderson et al. (2025).

**Table S16: Genes associated with climate change loci of interest**

**Table S17: Climatic niche parameters derived from climate-driven demographic model and used in Genome-Wide Association Study (GWAS)**

**Table S18: Climatic variables used in the genome-environment association (GEA)** analysis, including the source of the data, the code for each climatic factor, and the description of the climatic factor.

**Table S19: The GEA SNPs were associated with 37 climatic factors,** which can be assigned to five major groups: (a) snow dynamics (maximum snow depth, annual snow cover, annual snow duration, and the average, maximum and minimum timing of snowmelt), (b) total accumulated precipitation (annually and in all months except May), (c) the breadth of mean growing season temperature, (d) maximum temperature (annually, over the growing season and in all months except January), (e) and minimum temperature in five months (March, April, May, June, November). Many loci showed significant associations with multiple correlated climate variables. For example, SNP 1_21062864 was significantly associated with 17 climatic factors (three snow variables, 13 maximum temperature variables, and 2 minimum temperature variables). To avoid redundant analyses of genotype by climate interactions for highly correlated climatic factors, we examined the correlations of variables within each of the main five categories (snow dynamics, precipitation, mean temperature, maximum temperature, minimum temperature) to identify a single variable that reflects each of these axes of climatic variation. We found that the following five climatic factors were moderately to highly correlated with the other variables within each group: the timing of snowmelt, accumulated precipitation during the growing season, maximum growing season temperature, and minimum growing season temperature. These five tables show the correlations within climatic factors, beginning with correlation coefficients amongst snow dynamic traits, with correlation coefficients (r) and p-values. We have not corrected any of the p-values in these tables for multiple tests, as we were not specifically testing hypotheses with these models.

**Table S20: Species used to identify orthologues in OrthoFinder**, their source database, and accession information.
