## Supplemental Figures 1-11 for "The genetic basis of adaptation to continuous environmental gradients informs predictions about climate change responses"

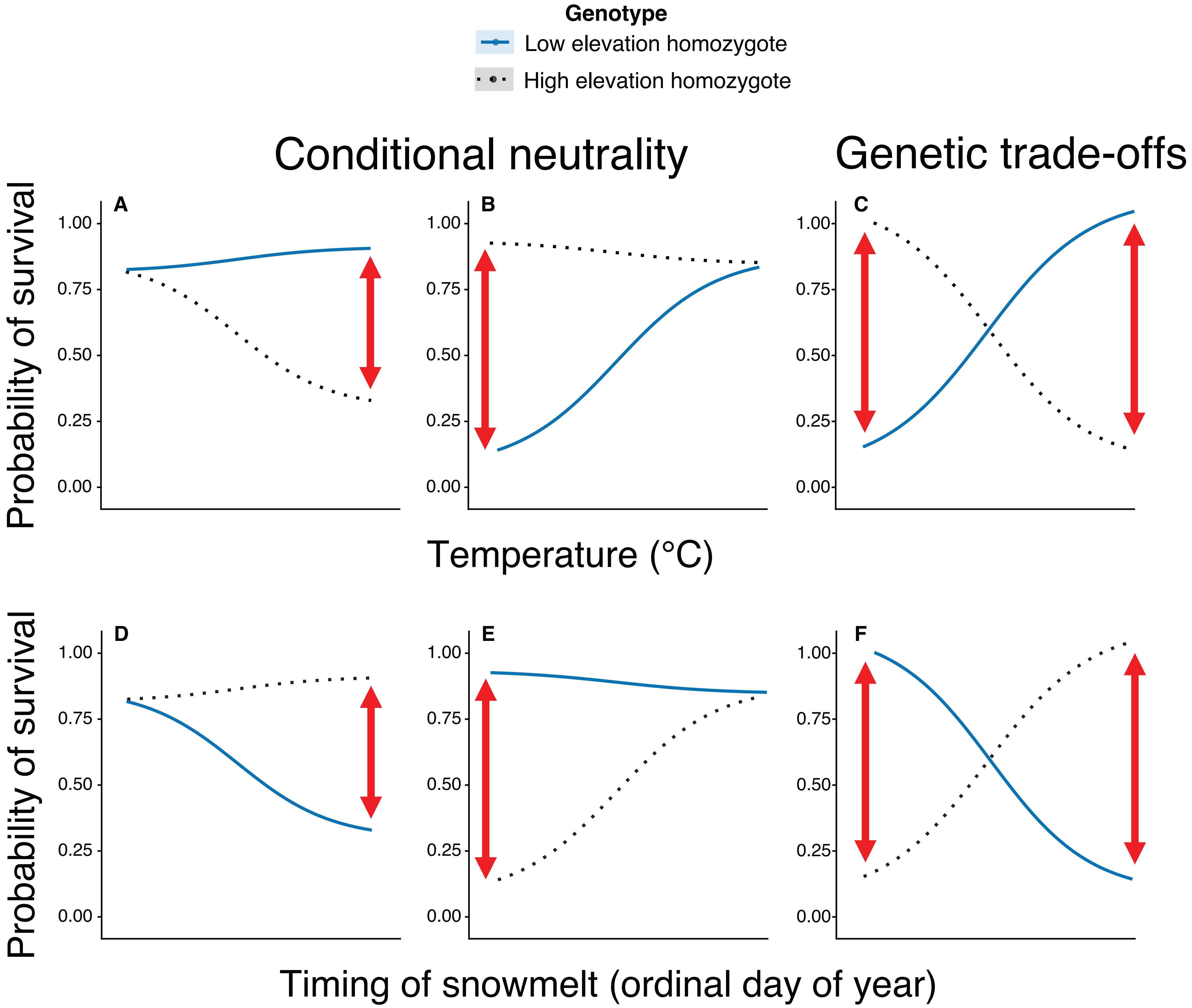

Genotype
Low elevation homozygote
High elevation homozygote
Conditional neutrality
Genetic trade-offs
A
B
C
D
E
F
Probability of survival
1.00
0.75
0.50
0.25
0.00
Temperature (°C)
Timing of snowmelt (ordinal day of year)

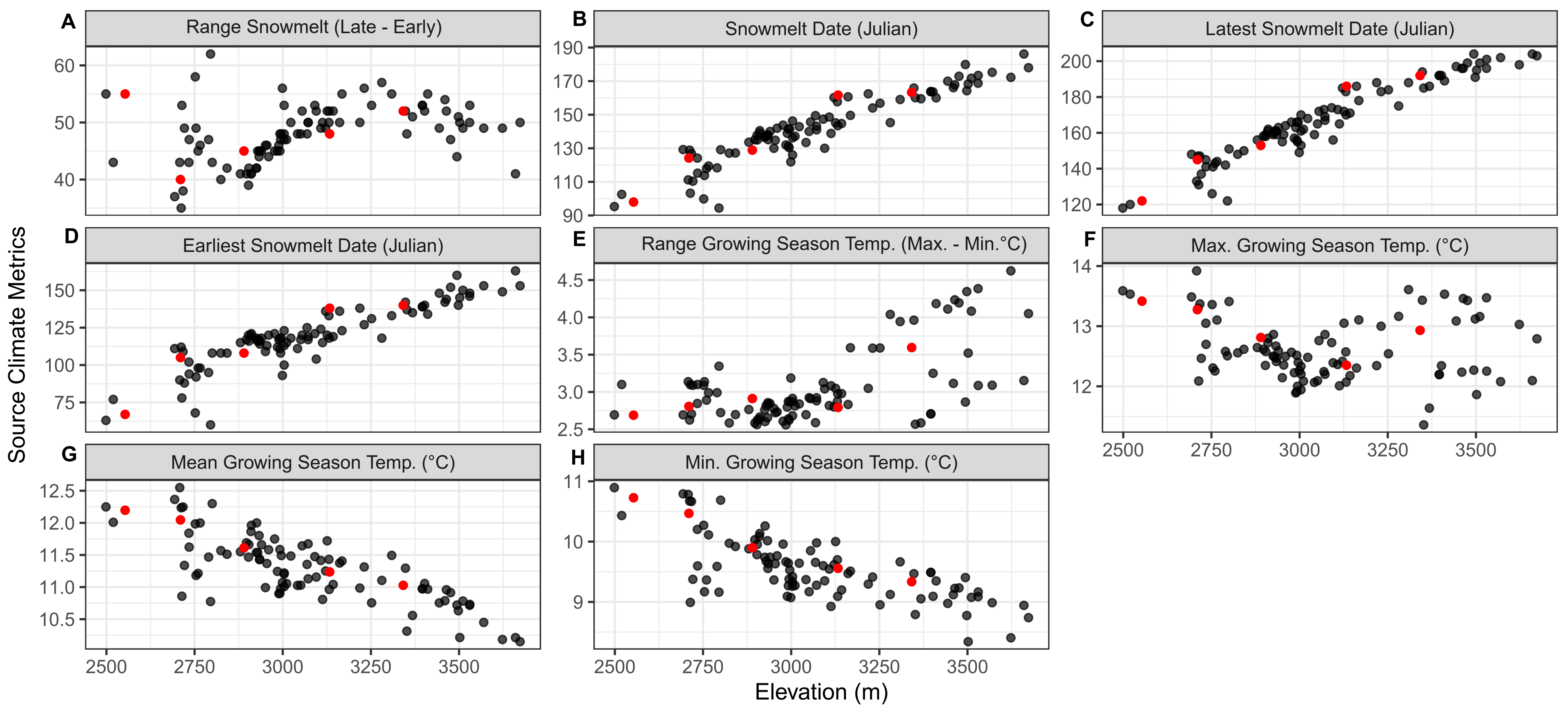

A
Range Snowmelt (Late - Early)
B
Snowmelt Date (Julian)
C
Latest Snowmelt Date (Julian)
D
Earliest Snowmelt Date (Julian)
E
Range Growing Season Temp. (Max. - Min.°C)
F
Max. Growing Season Temp. (°C)
G
Mean Growing Season Temp. (°C)
H
Min. Growing Season Temp. (°C)
Source Climate Metrics
Elevation (m)
2500
2750
3000
3250
3500

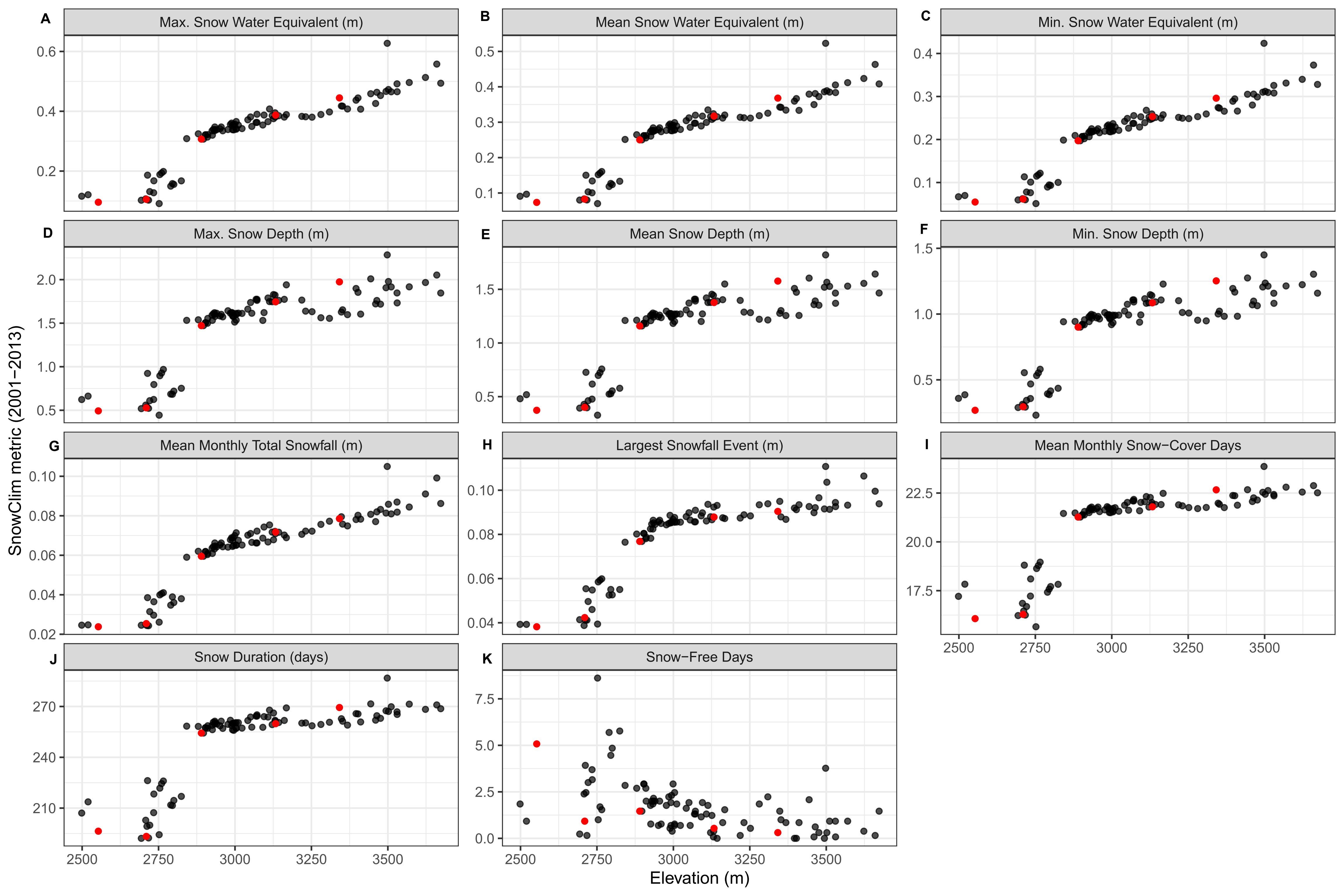
A
Max. Snow Water Equivalent (m)
B
Mean Snow Water Equivalent (m)
C
Min. Snow Water Equivalent (m)
D
Max. Snow Depth (m)
E
Mean Snow Depth (m)
F
Min. Snow Depth (m)
G
Mean Monthly Total Snowfall (m)
H
Largest Snowfall Event (m)
I
Mean Monthly Snow−Cover Days
J
Snow Duration (days)
K
Snow−Free Days
SnowClim metric (2001−2013)
Elevation (m)

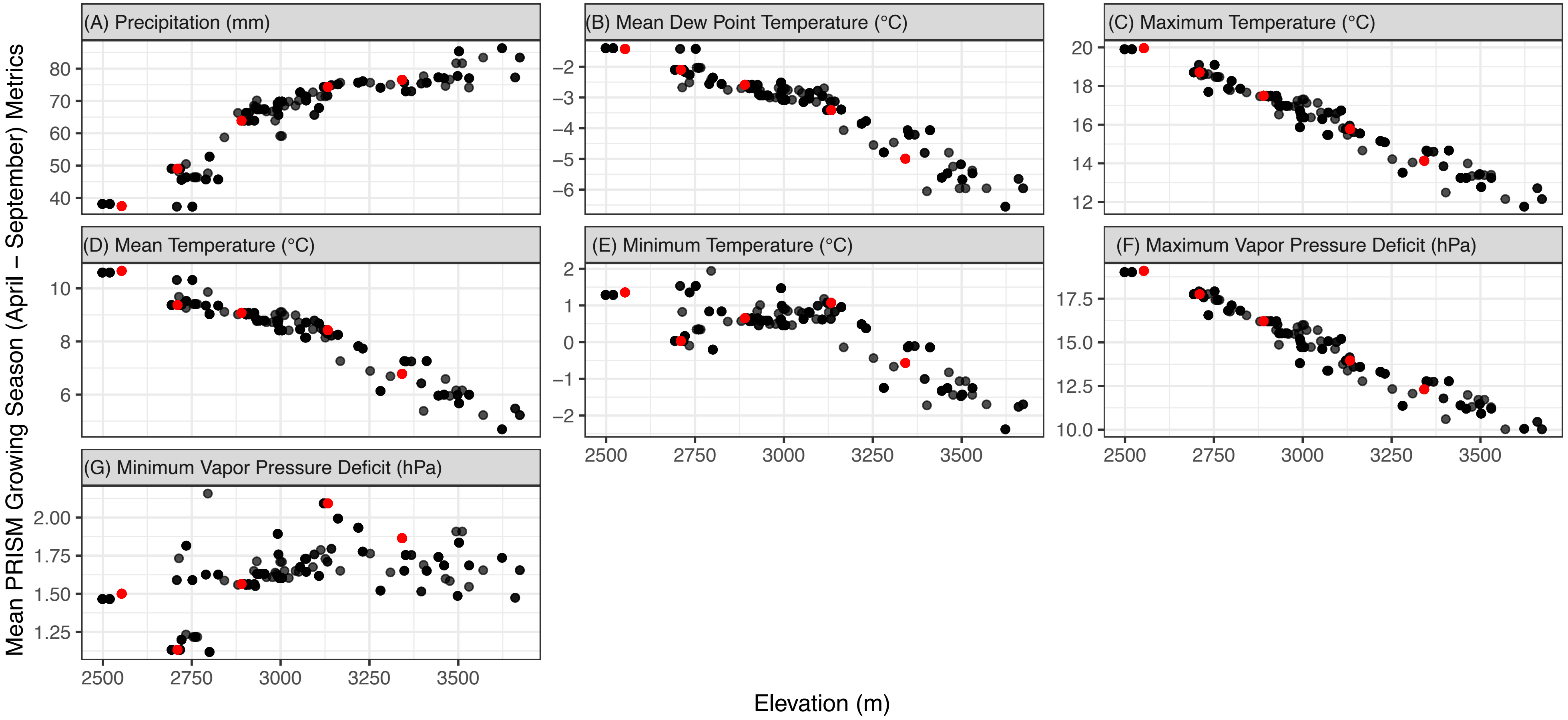

(A) Precipitation (mm)
(B) Mean Dew Point Temperature (°C)
(C) Maximum Temperature (°C)
(D) Mean Temperature (°C)
(E) Minimum Temperature (°C)
(F) Maximum Vapor Pressure Deficit (hPa)
(G) Minimum Vapor Pressure Deficit (hPa)
Mean PRISM Growing Season (April – September) Metrics
Elevation (m)
2500
2750
3000
3250
3500

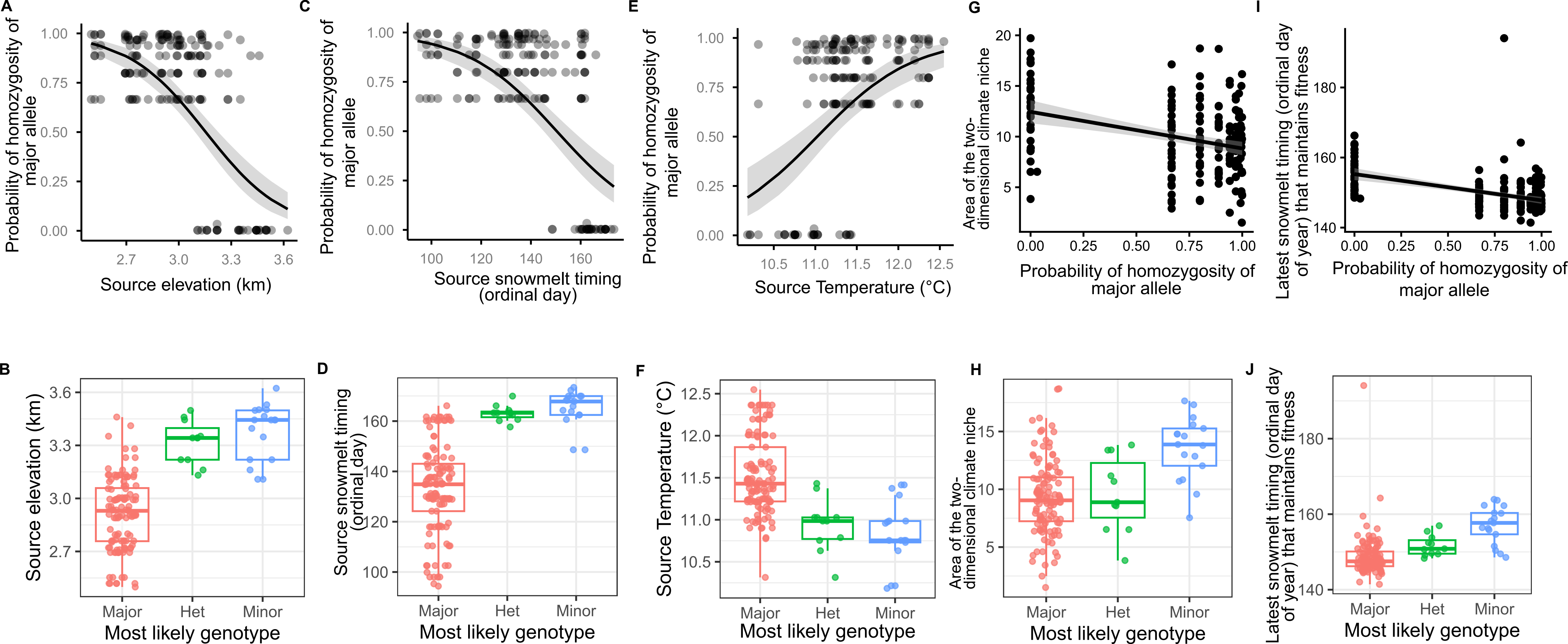

A
Probability of homozygosity of major allele
Source elevation (km)
C
Probability of homozygosity of major allele
Source snowmelt timing (ordinal day)
E
Probability of homozygosity of major allele
Source Temperature (°C)
G
Area of the two-dimensional climate niche
Probability of homozygosity of major allele
I
Latest snowmelt timing (ordinal day of year) that maintains fitness
Probability of homozygosity of major allele
B
Source elevation (km)
Major
Het
Minor
Most likely genotype
D
Source snowmelt timing (ordinal day)
Most likely genotype
F
Source Temperature (°C)
Most likely genotype
H
Area of the two-dimensional climate niche
Most likely genotype
J
Latest snowmelt timing (ordinal day of year) that maintains fitness
Most likely genotype

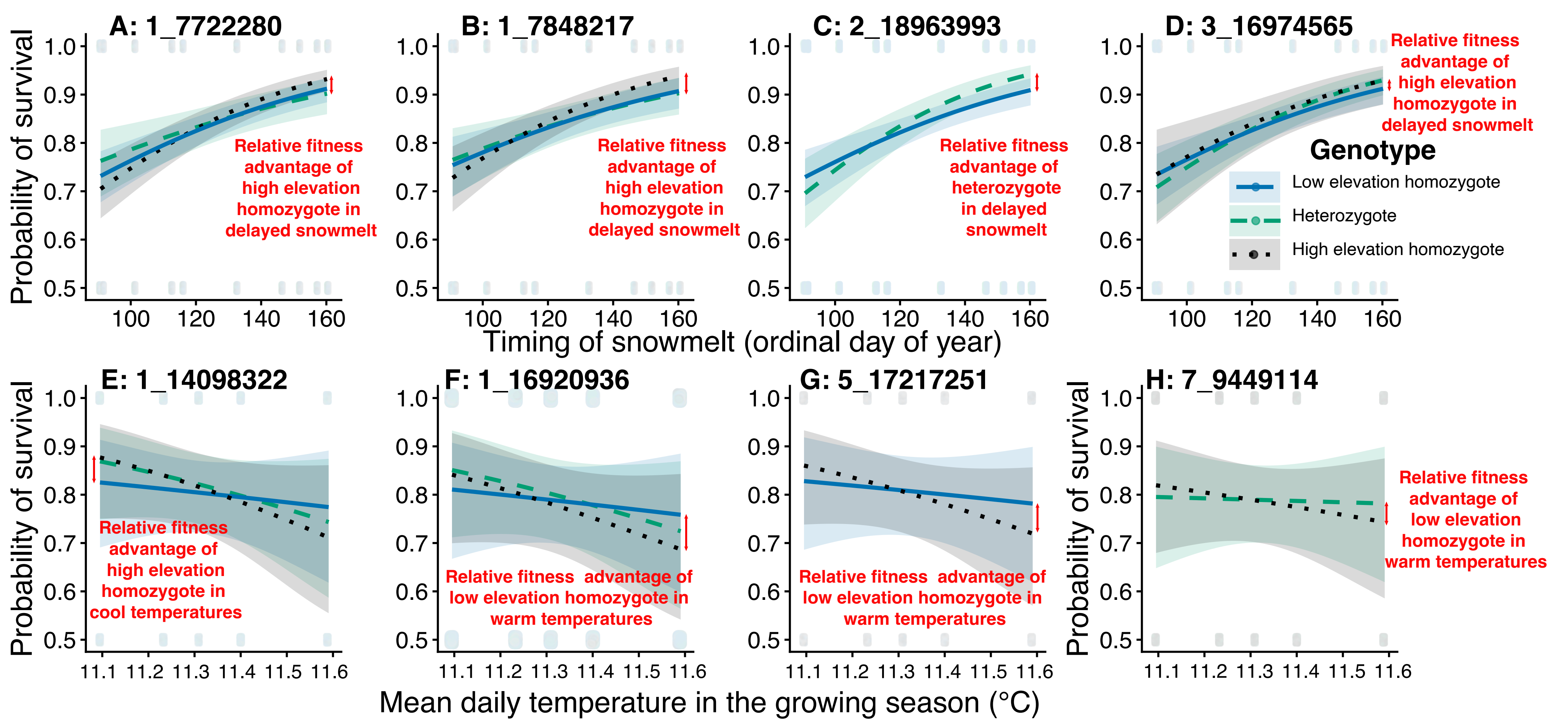
A: 1_7722280
B: 1_7848217
C: 2_18963993
D: 3_16974565
Probability of survival
Relative fitness advantage of high elevation homozygote in delayed snowmelt
Relative fitness advantage of high elevation homozygote in delayed snowmelt
Relative fitness advantage of heterozygote in delayed snowmelt
Relative fitness advantage of high elevation homozygote in delayed snowmelt
Genotype
Low elevation homozygote
Heterozygote
High elevation homozygote
Timing of snowmelt (ordinal day of year)
E: 1_14098322
F: 1_16920936
G: 5_17217251
H: 7_9449114
Relative fitness advantage of high elevation homozygote in cool temperatures
Relative fitness advantage of low elevation homozygote in warm temperatures
Relative fitness advantage of low elevation homozygote in warm temperatures
Relative fitness advantage of low elevation homozygote in warm temperatures
Mean daily temperature in the growing season (°C)

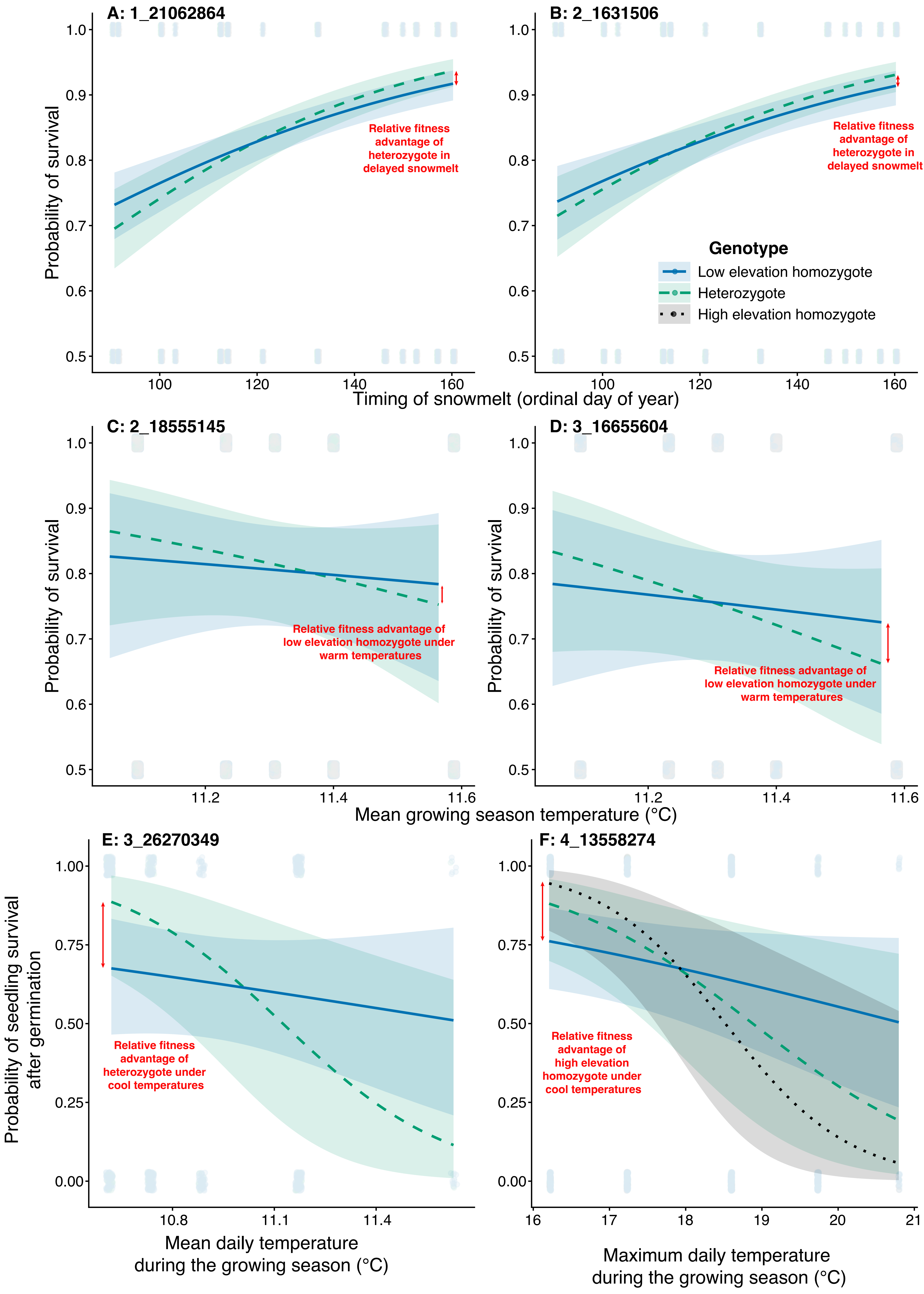
A: 1_21062864
B: 2_1631506
Probability of survival
Relative fitness advantage of heterozygote in delayed snowmelt
Genotype
Low elevation homozygote
Heterozygote
High elevation homozygote
Timing of snowmelt (ordinal day of year)
C: 2_18555145
D: 3_16655604
Relative fitness advantage of low elevation homozygote under warm temperatures
Mean growing season temperature (°C)
E: 3_26270349
F: 4_13558274
Probability of seedling survival after germination
Relative fitness advantage of heterozygote under cool temperatures
Relative fitness advantage of high elevation homozygote under cool temperatures
Mean daily temperature during the growing season (°C)
Maximum daily temperature during the growing season (°C)

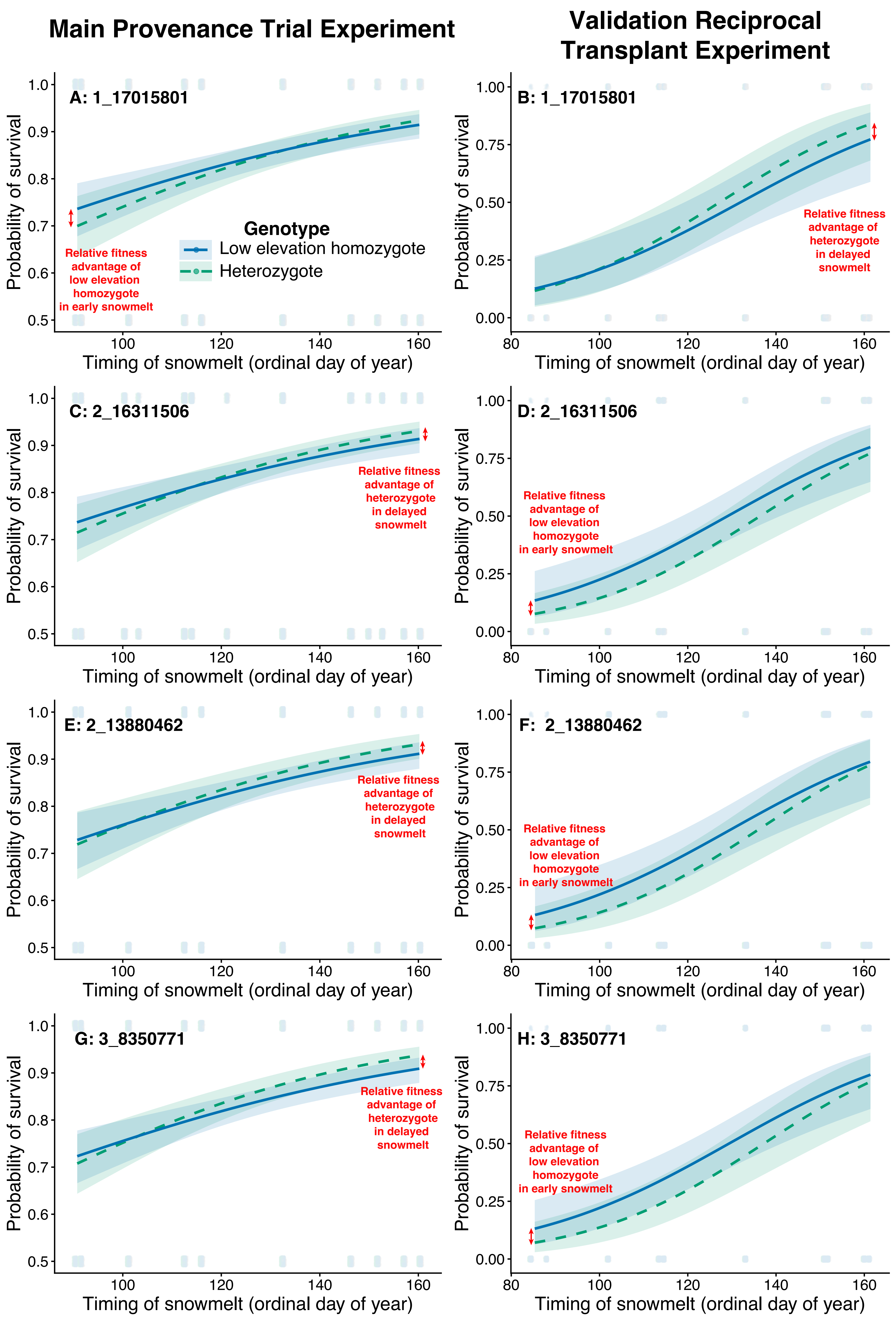
Main Provenance Trial Experiment
Validation Reciprocal Transplant Experiment
A: 1_17015801
B: 1_17015801
C: 2_16311506
D: 2_16311506
E: 2_13880462
F: 2_13880462
G: 3_8350771
H: 3_8350771
Genotype
Low elevation homozygote
Heterozygote
Relative fitness advantage of low elevation homozygote in early snowmelt
Relative fitness advantage of heterozygote in delayed snowmelt
Probability of survival
Timing of snowmelt (ordinal day of year)

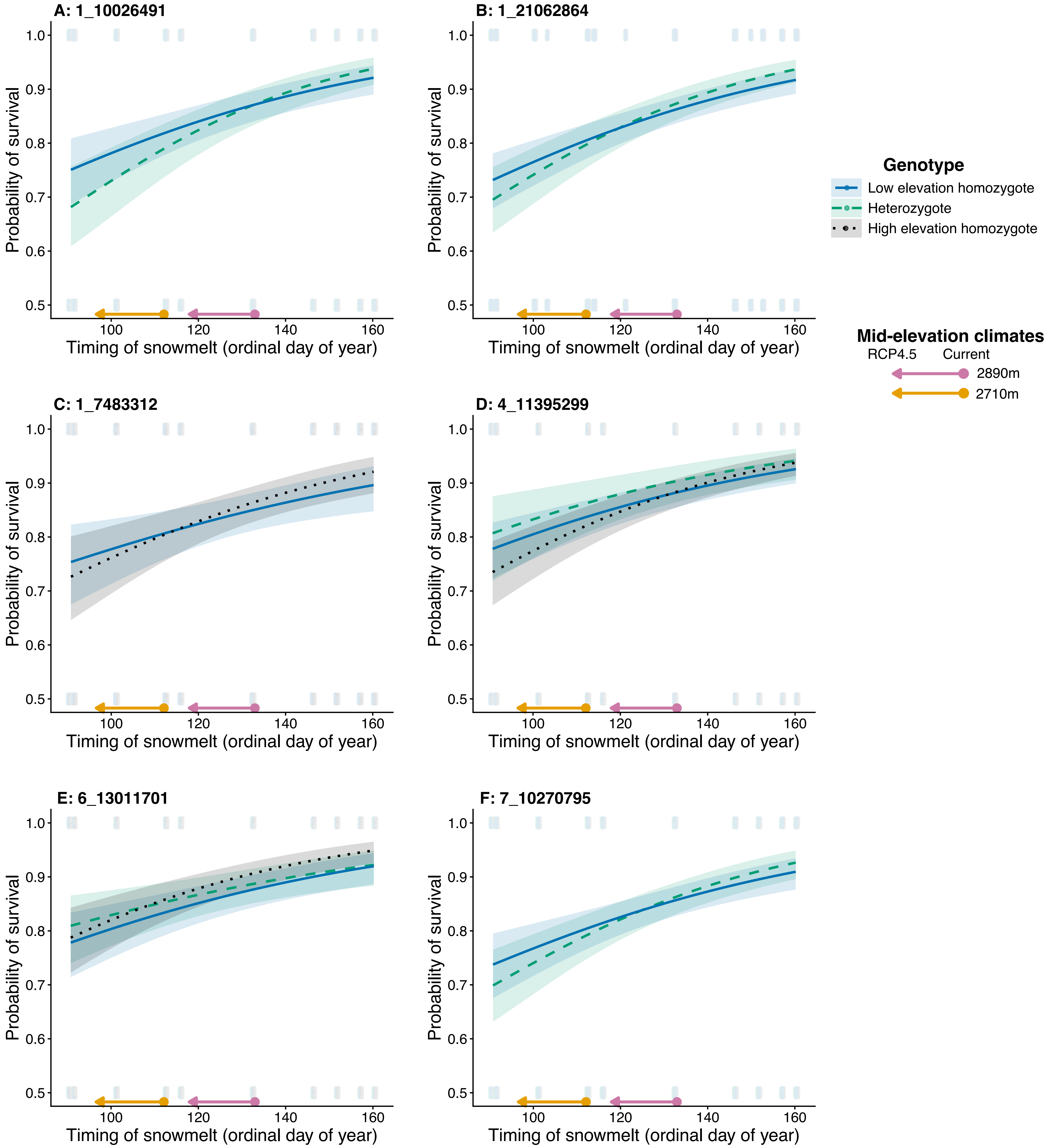
A: 1_10026491
B: 1_21062864
C: 1_7483312
D: 4_11395299
E: 6_13011701
F: 7_10270795
Probability of survival
Timing of snowmelt (ordinal day of year)
Genotype
Low elevation homozygote
Heterozygote
High elevation homozygote
Mid-elevation climates
RCP4.5
Current
2890m
2710m

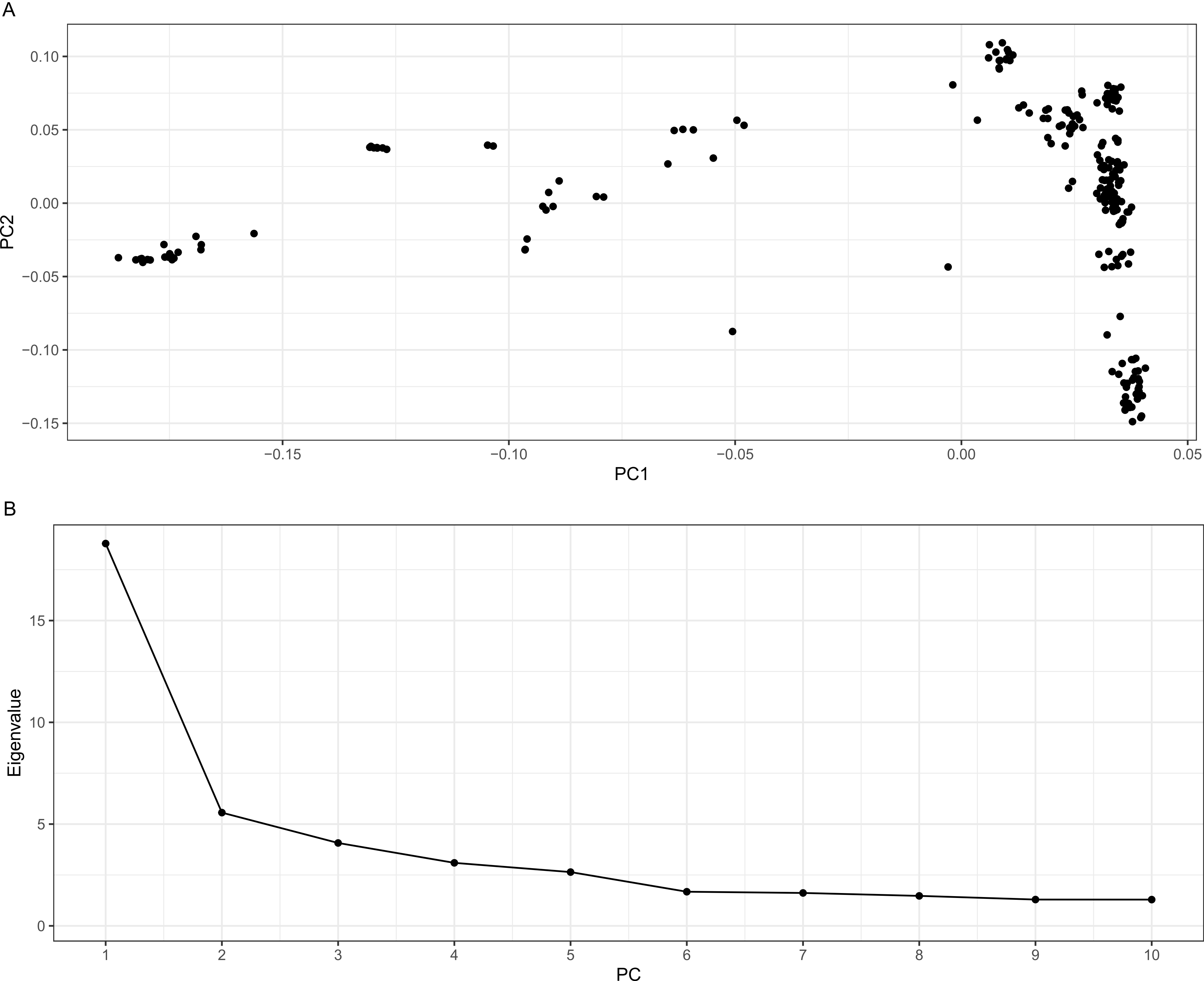

A
0.10
0.05
0.00
−0.05
−0.10
−0.15
PC2
−0.15
−0.10
−0.05
0.00
0.05
PC1
B
15
10
5
0
Eigenvalue
1
2
3
4
5
6
7
8
9
10
PC
